# A CCaMK/Cyclops response element in the promoter of *L. japonicus Calcium-Binding Protein 1* (*CBP1*) mediates transcriptional activation in root symbioses

**DOI:** 10.1101/2021.08.11.455944

**Authors:** Xiaoyun Gong, Elaine Jensen, Simone Bucerius, Martin Parniske

## Abstract

Early gene expression in arbuscular mycorrhiza (AM) and the nitrogen-fixing root nodule symbiosis (RNS) is governed by a shared regulatory complex. Yet many symbiosis-induced genes are specifically activated in only one of the two symbioses. The *Lotus japonicus* T-DNA insertion line T90, carrying a promoterless *uidA* (*GUS*) gene in the promoter of *Calcium Binding Protein1* (*CBP1*) is exceptional as it exhibits GUS activity in both root endosymbioses. To identify the responsible *cis-* and *trans*-acting factors, we subjected deletion/modification series of *CBP1* promoter:reporter fusions to transactivation and spatio-temporal expression analysis and screened EMS-mutagenized T90 populations for aberrant *GUS* expression. We identified one *cis-*regulatory element required for *GUS* expression in the epidermis and a second element, necessary and sufficient for transactivation by the Calcium and Calmodulin-dependent protein kinase (CCaMK) in combination with the transcription factor Cyclops and conferring gene expression during both AM and RNS. Lack of *GUS* expression in T90 *white* mutants could be traced to DNA hypermethylation detected in and around this element. We concluded that the CCaMK/Cyclops complex can contribute to at least three distinct gene expression patterns on its direct target promoters *NIN* (RNS), *RAM1* (AM), and *CBP1* (AM and RNS), calling for yet-to-be identified specificity-conferring factors.

## Introduction

As main constituents of nucleotides, proteins and nucleic acids, nitrogen and phosphate are essential for life (Bowler *et al.*, 2010). Two types of plant root endosymbiosis, the arbuscular mycorrhiza (AM) and the nitrogen-fixing root nodule symbiosis (RNS) hold promise for sustainable agriculture. AM and RNS greatly benefit plant nutrition by improving nutrient uptake from the soil and providing ammonium as a nitrogen source, respectively. RNS can significantly reduce the demand for chemical nitrogen fertiliser application, hence reducing fossil fuel consumption and, once globally adjusted to a sustainable scale, the negative ecological impact imposed by the release of ammonium and nitrogen oxides into the atmosphere, groundwater, rivers, lakes and the sea (Fowler *et al.*, 2013).

Establishment of both AM and RNS requires chemical communications between symbiotic partners that induce concerted structural modification and rearrangement of the host cells, both controlled by a cohort of transcriptional circuitries. To unravel the genes underlying the development of these two important symbioses, transcriptome analysis has been employed and led to a catalogue of symbiosis-regulated genes that are responsive to AM (Liu *et al.*, 2003; Kistner *et al.*, 2005; Hogekamp *et al.*, 2011; Gutjahr *et al.*, 2015) or RNS (Demina *et al.*, 2013; Breakspear *et al.*, 2014; Roux *et al.*, 2014) . Although many symbiosis regulated genes are expressed specifically in either RNS or AM, such as *Nodule Inception* (*NIN*) (Schauser *et al.*, 1999; Kumar *et al.*, 2020) or *Reduced Arbuscular Mycorrhiza 1* (*RAM1*) (Gobbato *et al.*, 2012), respectively, a small subset of genes appears to be induced in both symbioses. These include a symbiosis induced Subtilase (*SbtS*) (Kistner *et al.*, 2005; Takeda *et al.*, 2007, 2009), *Vesicle-Associated Membrane Protein 72* (Ivanov *et al.*, 2012), *ENOD11* (Boisson-Dernier *et al.*, 2005)*, Vapyrin* (Murray *et al.*, 2011) and *ABC-B transporters in Mycorrhization and Nodulation* genes (Roy *et al.*, 2021).

The question how the three different patterns of gene expression in response to symbiotic bacteria and fungi are accomplished, is particularly puzzling, because early gene expression in both symbioses depends on the same subset of the so called “common symbiosis genes”, some of which encode proteins involved in early signal transduction processes (Kistner *et al.*, 2005; Oldroyd, 2013). A key to specificity may be at the initiation step of the signalling cascade because microsymbiont-derived molecules, lipo-chitooligosaccharides (LCOs) with specificity-conferring decorations produced by nitrogen-fixing bacteria (collectively referred to as Nod factor), or LCOs and short chain chitin oligosaccharides (COs) by AM fungi (Maillet *et al.*, 2011; Radutoiu *et al.*, 2019) are believed to be perceived by distinct complexes comprising LysM type receptor-like kinases (Radutoiu *et al.*, 2003; Zhang *et al.*, 2021). Symbiosis Receptor-like Kinase (SymRK) (Endre *et al.*, 2002; Stracke *et al.*, 2002) can associate with specific LysM receptors and thus forms a conceptual link between the perception of microbial (L)COs and the initiation of symbiotic downstream responses (Ried *et al.*, 2014; Antolín-Llovera *et al.*, 2014). A hallmark of the common signalling process is the generation of nuclear calcium oscillation (or spiking) (Sieberer *et al.*, 2009) facilitated by ion channels and transporters on the nuclear envelope (see review by Kim et al., 2019). Calcium spiking is postulated to act as a second messenger which is presumably decoded in the nucleus by a Calcium-Calmodulin dependent kinase (CCaMK) (Lévy *et al.*, 2004; Tirichine *et al.*, 2006). CCaMK is activated *via* binding of a, yet to be identified, calmodulin, and interacts with and phosphorylates Cyclops, a DNA-binding transcription factor (Yano *et al.*, 2008; Miller *et al.*, 2013; Singh *et al.*, 2014). This protein complex is required for both RNS and AM to activate symbiosis-related genes, e.g. *RAM1* during AM (Pimprikar *et al.*, 2016); or *NIN* and *ERF Required for Nodulation 1* (*ERN1*) during RNS (Singh *et al.*, 2014; Cerri *et al.*, 2017).

*Cis-*acting regulatory sequences (*cis*-elements) are crucial for temporal and/or spatial regulation of gene expression in eukaryotes. In agreement with this, RNS or AM-related *cis-* elements have been identified in the promotors or intronic regions of symbiosis-regulated genes (Pimprikar & Gutjahr, 2018; Liu *et al.*, 2019; Akamatsu *et al.*, 2020). Taking into account the three principally different expression patterns (AM-induced, RNS-induced and commonly induced) in the light of the postulate of a common symbiosis signalling pathway, it was hypothesised that the different gene expression patterns are achieved by two independent and symbiosis-specific pathways that act in parallel to the common signalling pathway (Schultze & Kondorosi, 1998). According to this model, dual gene expression in both symbioses could be achieved by promoters harbouring both AM-specific and RNS-specific *cis*-elements, thus accumulating the output of the two specific and independent pathways, or *cis*-elements exclusively responsive to the output of the common symbiosis pathway, or a mix of all three types of *cis*-elements.

To obtain further insights into the mechanisms that confer common symbiosis-related gene expression, we employed the *Lotus japonicus* promoter tagging line T90 (Webb *et al.*, 2000), which has served as a useful marker line for the study of plant symbiotic signal transduction over the last two decades (Kistner *et al.*, 2005; Gossmann *et al.*, 2012; Ried *et al.*, 2014; Banhara *et al.*, 2015). T90 carries a single copy of a T-DNA, containing a promoterless *GUS* gene, which is inserted in the promoter region of the *Calcium Binding Protein 1* gene (*CBP1*; Fig. **1a**). The T90 *GUS* gene expression was so far exclusively observed in plant roots inoculated with AM fungi (Kistner *et al.*, 2005) or rhizobia, including *Mesorhizobium loti* strain R7A in an NF-dependent manner (Webb *et al.*, 2000) and treated with *M. loti* strain R7A Nod factor (Gossmann *et al.*, 2012; Webb *et al.*, 2000; Fig.**1b** & **S1**) but in no other tissues or treatments tested (Kistner *et al.*, 2005; Gossmann *et al.*, 2012). For example, T90 *GUS* expression was neither detected in T90 shoots or leaves nor inducible by synthetic hormones 1-Naphthaleneacetic acid (NAA) or 6-Benzylaminopurine (6-BAP) (Webb *et al*., 2000; Tuck, 2006). It was also not induced upon inoculation with the growth-promoting fungus *Serendipita indica* (previously known as *Piriformospora indica*) (Banhara *et al*., 2015).

**Fig. 1.**
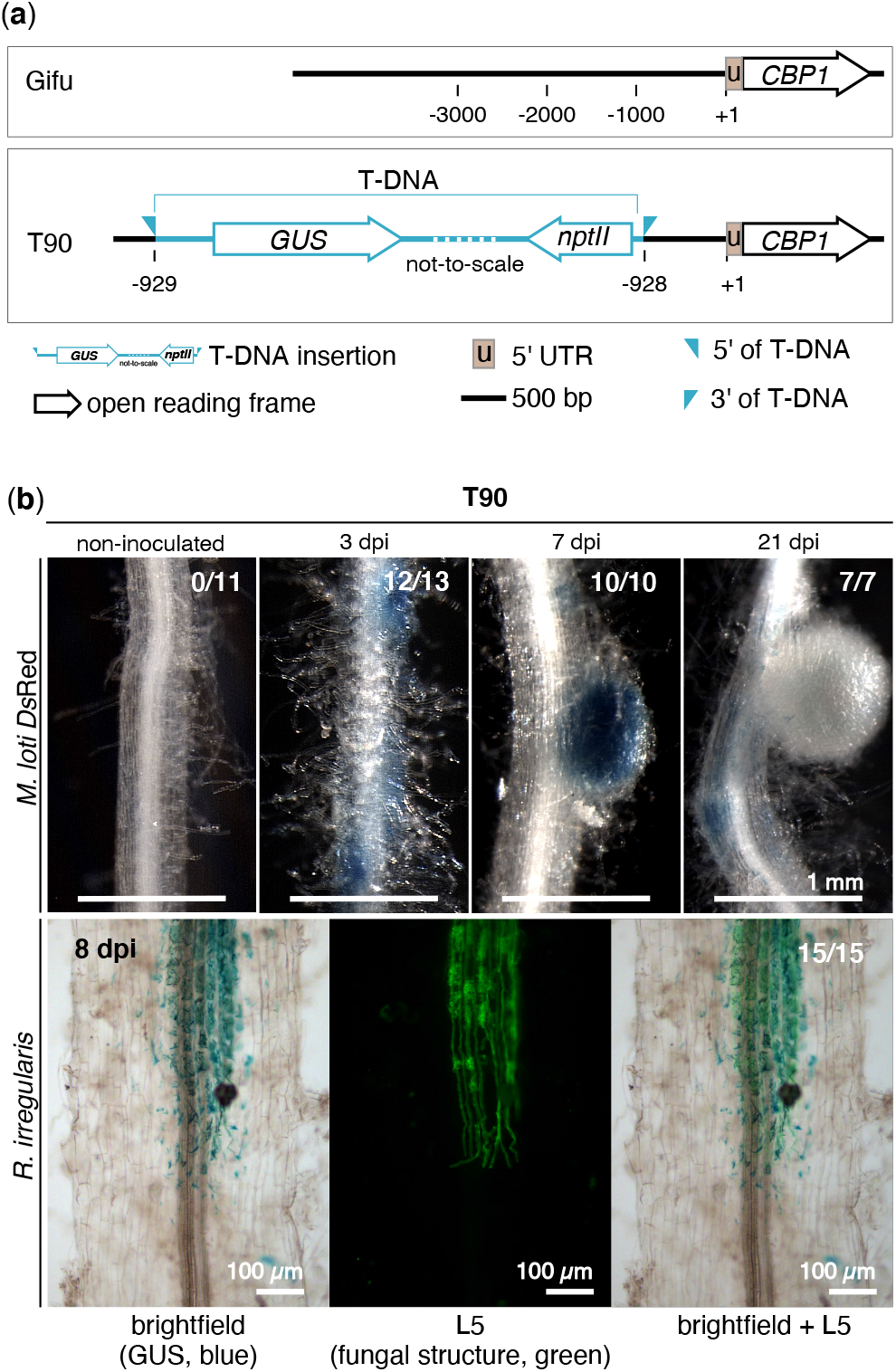
*GUS* expression pattern in T90 roots. (a) Position of the T-DNA insertion within the promoter of *CBP1* gene in the transgenic line T90. The respective region in *L. japonicus* ecotype Gifu is shown as a reference. (b) T90 roots were stained with X-Gluc to reveal a blue coloration generated by GUS enzyme activity at indicated days post inoculation (dpi) with *M. loti Ds* Red or AM fungus *Rhizophagus irregularis*. Note the blue staining at 3 dpi in patches in root hair cells, the presence and absence of blue staining in the central nodule tissue at 7 dpi and 21 dpi, respectively. Green: Alexa Fluor-488 WGA-stained *R. irregularis* visualised with a Leica Filter Cube L5. #/# place top right corner in (b): number of plants displaying GUS activity / total number of plants analysed. Sections of nodules or roots are displayed in Fig. S1.

This work aimed to decipher the molecular secret behind the common, yet exclusive symbiosis-induced gene expression pattern of T90. To this end, we performed a classical “forward genetics” approach in which we generated an ethyl methanesulfonate-mutagenised T90 population and screened M_2_ and M_3_ families for the loss and gain of *GUS* expression. In parallel, we used promoter deletion series to identify regions with relevance for symbiotic responsiveness.

## Material and Methods

**Plant, bacterial and fungal material** is listed in Method S1.

**Plant growth conditions and phenotypic analysis** for plants shown in each figure is included in Method S2.

### Promoter analysis and microscopy

*Lotus japonicus* hairy roots were generated using an *Agrobacterium rhizogenes* based protocol (Charpentier *et al.*, 2008) with the following modifications: 1) roots of seedlings were cut away while seedlings were immersed in *A. rhizogenes* that was re-suspended in sterile MilliQ water; 2) after removal of roots, shoots of the seedlings were transferred to plates containing Gamborg’s B5 medium (without sucrose) and 0.8 % BD Bacto™ agar. The T-DNA region of the construct carried in *A. rhizogenes* contained a GFP transformation marker (*Ubi10_pro_:NLS-2xGFP*) and a promoter:GUS reporter fusion placed in tandem. Chimeric root systems with transformed roots and/or nodules were identified by GFP fluorescence emanating from nuclei under a Leica MZ16 FA fluorescent stereomicroscope equipped with a GFP3 filter (Leica Microsystems). Transformed nodule primordia and nodules were identified by the presence of a red fluorescence signal under a *Ds*Red filter 10 to 14 dpi, then excised and subject to GUS staining for 3 h at 37 ^o^C (Groth *et al.*, 2010 with incubating time adjusted). The whole root systems were subjected to GUS staining for 6 h to investigate the GUS activity across the whole roots (Fig. **3**,**S3,S5**). For promoter analysis of mycorrhized roots, roots were subject to GUS staining for 14 to 16 h at 37 °C, followed by staining with WGA (Method S3). Microscopic procedure is detailed in Method S4.

**Fig. 2.**
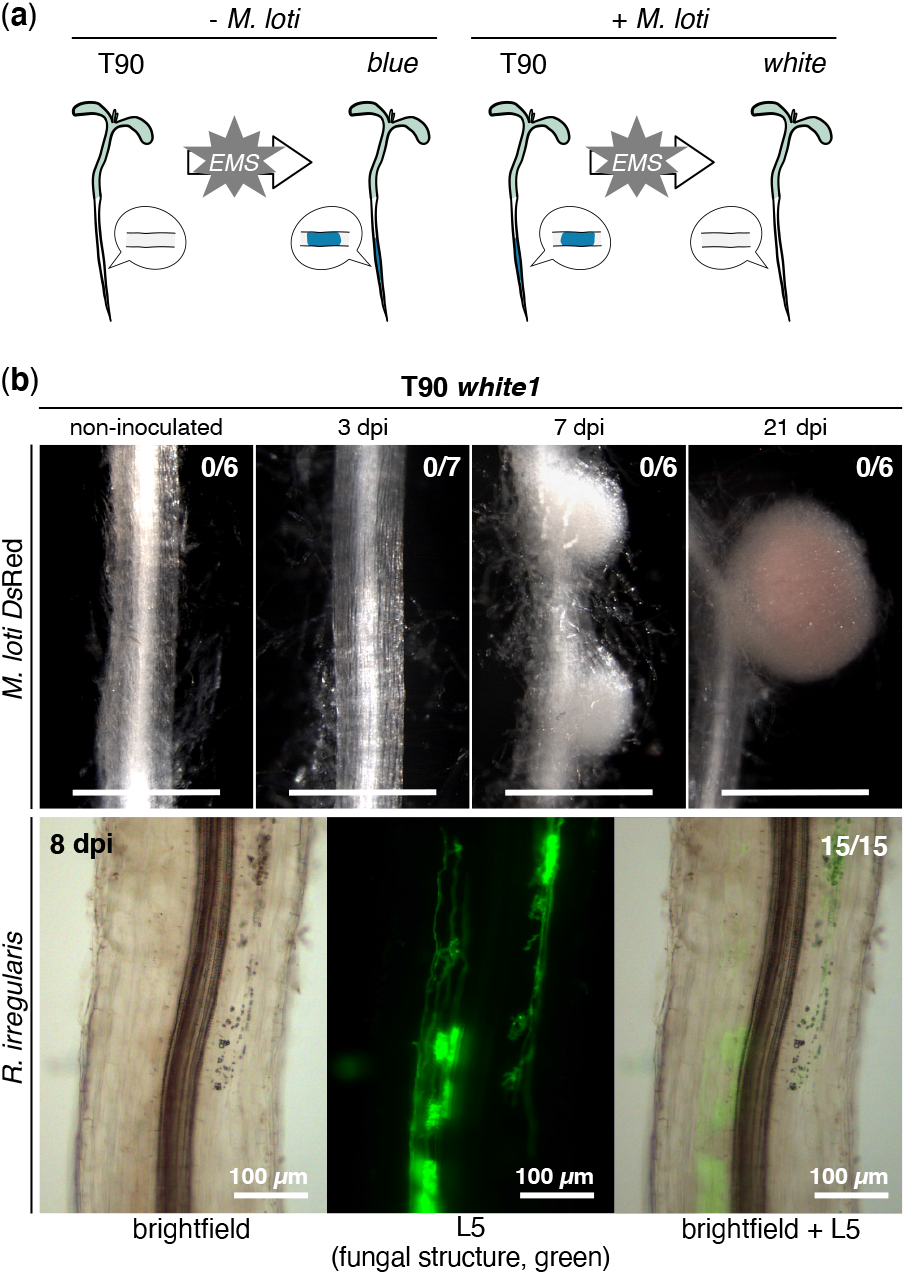
Absence of GUS activity in T90 *white* mutant roots during AM or RNS. (a) Schematics of the two screens of EMS- induced mutant populations for M_2_ seedlings with altered GUS activity: spontaneous activation of the *GUS* gene in the absence of symbionts (left) or undetectable GUS activity in the presence of symbionts (right; resulting mutants are referred to as T90 *white* mutants). (b) T90 *white1* roots were stained with X-Gluc to reveal GUS activity at indicated dpi with *M. loti Ds*Red or *R. irregularis*. Note the total absence of GUS activity in T90 *white* roots, compared to those of T90 upon inoculation with microsymbionts (tested side-by-side in the same experiment; see Fig. 1a; Fig. S1). Pictures of T90 *white1* root systems and analysis of T90 *white3* are included in Fig. S2c-d. Green: Alexa Fluor-488 WGA-stained *R. irregularis* visualised with a Leica Filter Cube L5. #/# top right corner of images: number of plants displaying GUS activity / total number of plants analysed.

**Fig. 3.**
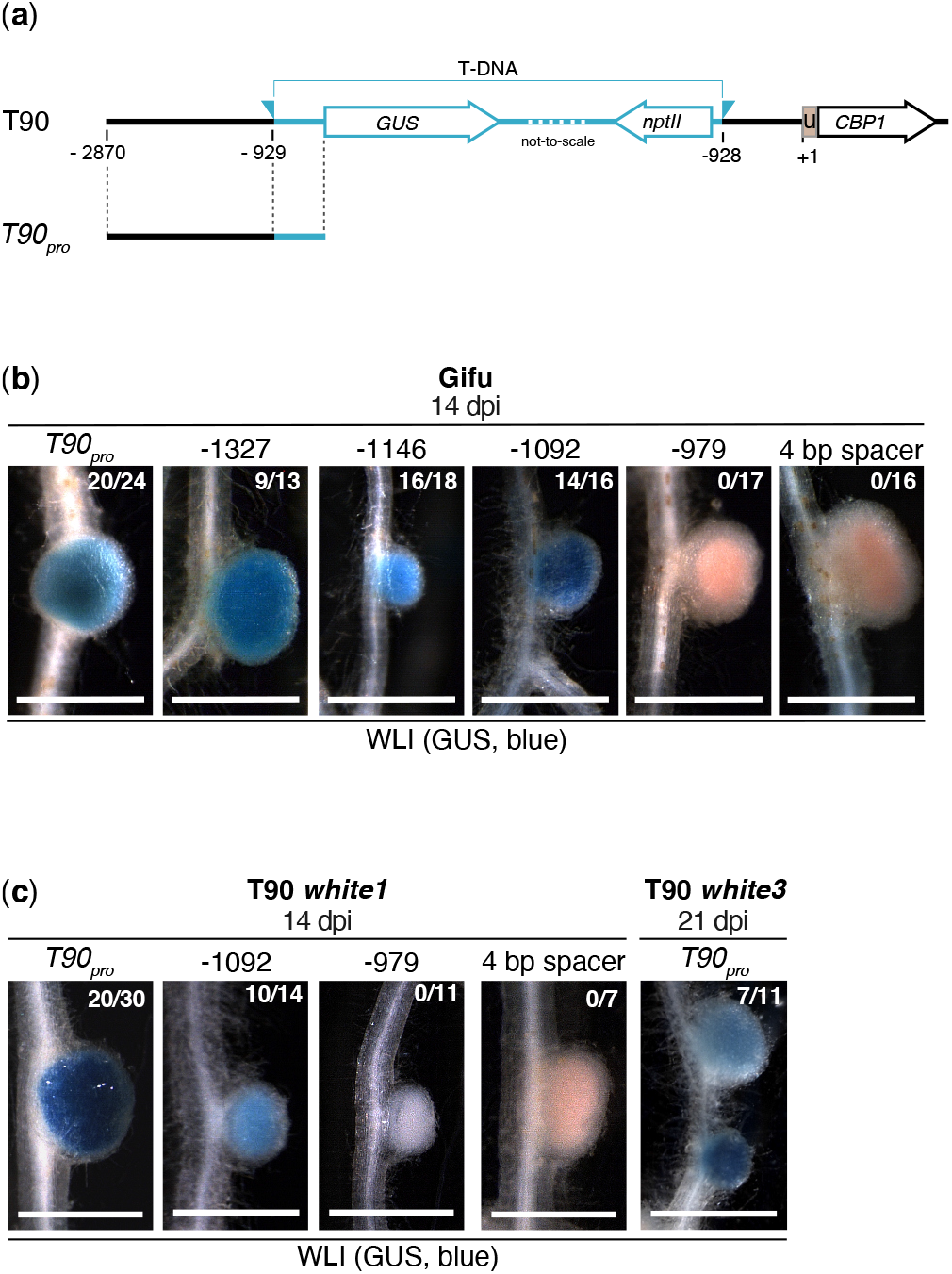
Absence of GUS activity in T90 *white* mutants can be restored by transgenic *T90_pro_:GUS* fusions. (a) T-DNA insertion in the T90 line as in Fig. 1, here including the *CBP1* promoter reference coordinates used for definition of the length of the individual fragments tested in the T90 promoter deletion series (for symbols refer to Fig. 1b). (b) *L. japonicus* Gifu or (c) T90 *white* mutants (*white1* or *white3*) hairy roots transformed with T-DNAs carrying a *Ubq10_pro_:NLS-GFP* transformation marker together with a *GUS* reporter gene driven by the full length T90 promoter (−2870) or a deletion series thereof (starting at −1327, −1146, −1092 or −979 of the *CBP1* promoter), or a 4 bp spacer sequence were analysed at indicated with *M. loti Ds*Red. Pictures of entire root systems from these experiments are included in Fig. S3a-b. #/#: number of plants showing GUS activity in nodules / total number of transgenic root systems analysed. BF: brightfield. Bars, 1 mm unless stated otherwise. WLI: white light illumination.

### DNA constructs

A detailed description of the constructs used in this study is provided in Table S2. Constructs were generated with the Golden Gate cloning system described in Binder *et al. (*2014). A variant of the *GUS* gene, *DoGUS* (from plasmid C204, DNA Cloning Service), adapted for the Golden Gate cloning system was kindly provided by David Chiasson (SMU, Halifax, Canada).

### *EMS* mutagenized T90 population and mutant screening

Generation of the ethyl methane sulphonate (EMS) mutagenized T90 population is described in the *Lotus japonicus* handbook section ‘*EMS mutagenesis*’ (Márquez, 2005). Details of how plants were grown and screening was carried out are listed in Method S5.

### Transient expression assays in *Nicotiana benthamiana* leaves

*N. benthamiana* plants were grown as previously described (Cerri *et al.*, 2017), and infiltration of *N. benthamiana* leaves with *A. tumefaciens* was performed as previously described (Cerri *et al.*, 2012) but with acetosyringone concentration modified to 150 µM. *A. tumefaciens* carrying *promoter:GUS* fusion constructs of interest (strain AGL1) were co-infiltrated with *A. tumefaciens* containing plasmid *35S_pro_:3xHA-Cyclops* (strain AGL1; Singh *et al.*, 2014), *35S_pro_:CCaMK^1-314^-NLS-mOrange* (strain GV3101; Takeda *et al.*, 2012) or *35S_pro_:CCaMK^T265D^-3xHA* (strain GV3101; Banba *et al.*, 2008) as indicated in Fig. **5** & **S3**. An AGL1 strain carrying a K9 plasmid constitutively expressing *Discosoma sp.* red fluorescent protein (*Ds*Red) was used as needed to equalize the amount of *A. tumefaciens* infiltrated per leaf, together with an *A. tumefaciens* strain carrying a plasmid for the expression of the viral P19 silencing suppressor to reduce post-transcriptional gene silencing (Voinnet *et al.*, 2003). *N. benthamiana* leaf discs with a diameter of 0.5 cm were harvested at 60 hours post infiltration. A total number of 8 leaf discs per indicated vector combination were analysed in at least 2 independent experiments performed in different weeks.

**Fig. 4.**
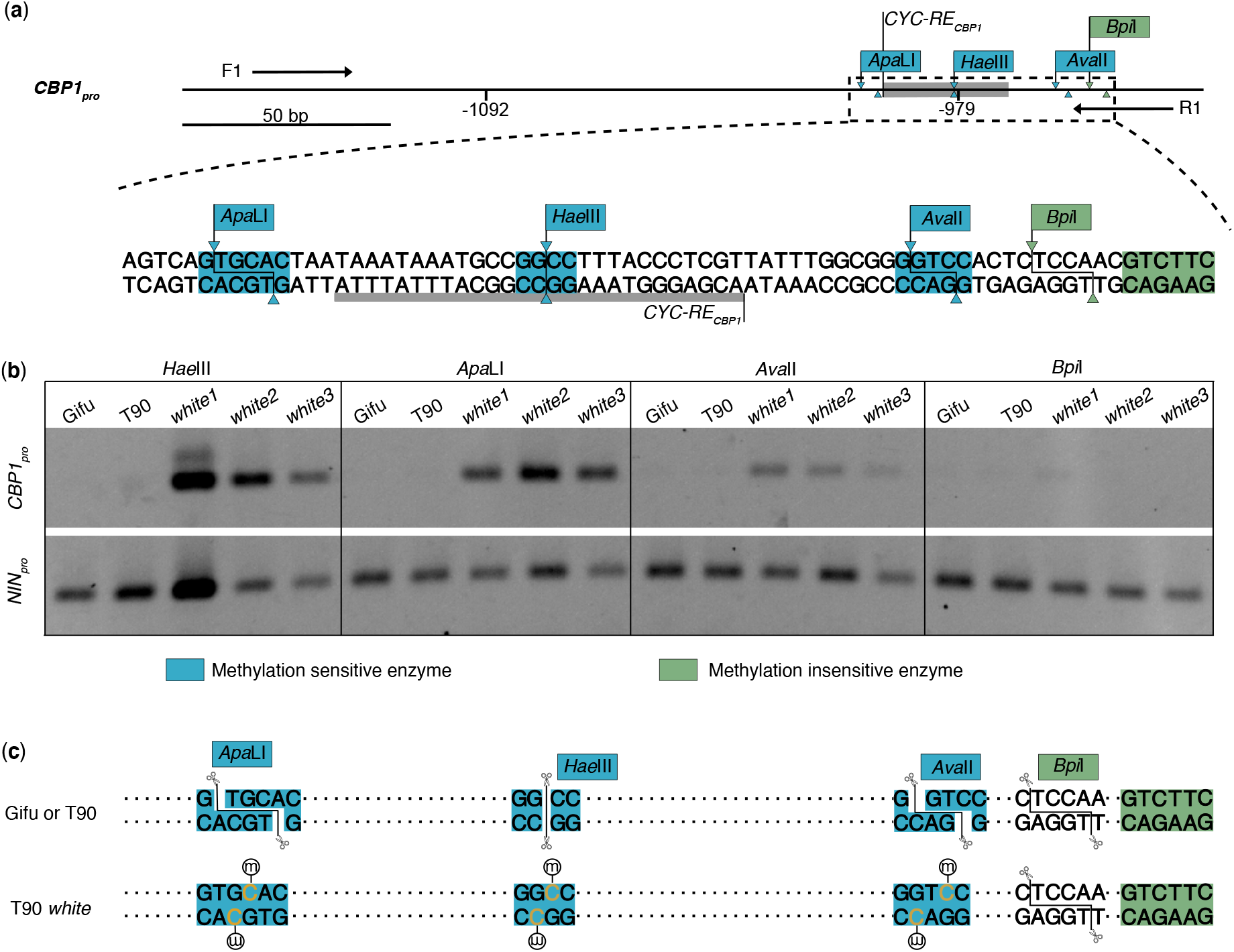
Cytosine methylation within and around a 113 bp T90 promoter region of T90 *white* mutants but not those of T90 or *L. japonicus* Gifu. (a) Genomic DNA (gDNA) from *L. japonicus* Gifu, T90 or T90 *white* mutants (*white1*, *white2* or *white3*) was digested with either one of the restriction digestion enzymes: methylation-sensitive restriction enzymes *Hae*III, *Apa*LI and *Ava*II (blue) or methylation-insensitive restriction enzyme *Bpi*I (green). Blue and green shades on DNA sequence: recognition sites of restriction enzymes. Grey shade in the restriction map or grey underline of promoter sequence: *CYC-RE_CBP1_.* Arrowheads and lines: endonucleolytic cleavage site and outline of the restriction digestion products. (b) Analysis of the success of restriction digestion by PCR. Digested gDNA from the indicated genotype was used as a template for PCR amplification with primers F1 and R1 flanking a 262 bp promoter region (top panel) and primers flanking a 199 bp stretch of the *L. japonicus NIN* promoter as control (bottom panel). This region in the *NIN* promoter does not contain restriction sites for any of the restriction enzymes used. This control indicates that the status of the gDNA was suitable for PCR after the digestion process. Note that PCR products could be obtained using digested gDNA from the T90 *white* mutants but not from Gifu or T90 as amplification template. (c) Graphic summary of the results in (b) projected onto the promoter region together with the recognition sites of the restriction enzymes. m in an open circle: methyl groups. Scissor cartoon: successful endonucleolytic cleavage.

**Fig. 5.**
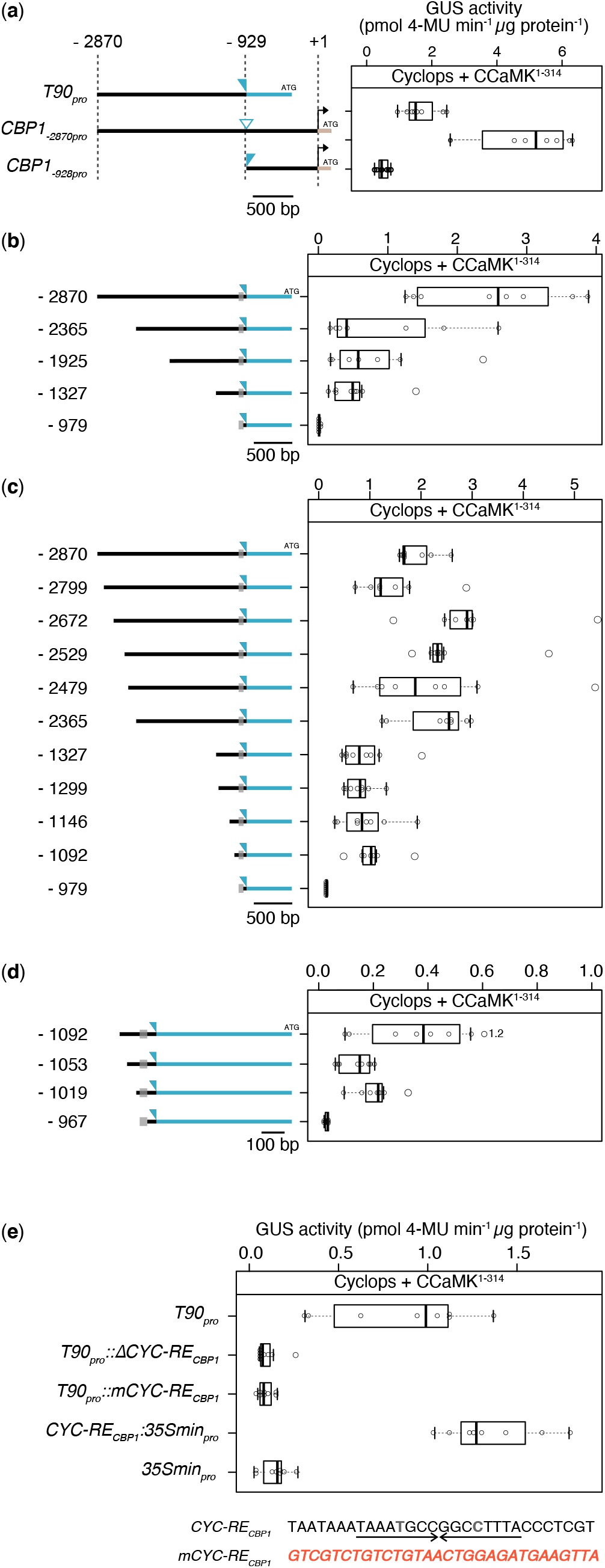
*A cis-*element in the promoter of *CBP1* is necessary and sufficient for the CCaMK^1-314^/Cyclops mediated transactivation of promoter:GUS reporter fusions in *Nicotiana benthamiana* leaf cells. (a-e) *N. benthamiana* leaf cells were transformed with T-DNAs carrying a *GUS* reporter gene driven by either of the indicated promoters: (a) the T90 promoter (labelled as *T90_pro_*in a & d or −2870 in b-c); one of the two *CBP1* promoter regions (*CBP1_-2870pro_*or *CBP1_-928pro_*); (b-d) promoter deletion series generated in the context of *T90_pro_* including promoter regions that were (b) ca. 300 - 500 bp different in length, ca. 50 to 100 bp different in length within −2870 to −2025 bp or −1327 to −979 bp, (d) ca. 35 - 50 bp different in length within −1092 to −967 bp; (e) *T90_pro_*, *T90_pro_* with the 30 nt long *cis-* element (*CYC-RE_CBP1_*) mutated or deleted (*T90_pro_::mCYC-RE_CBP1_* or *T90_pro_::∆CYC-RE_CBP1_*, respectively), a 35S minimal promoter (*35Smin_pro_*), or *CYC-RE_CBP1_*fused to *35Smin_pro_*(*CYC-RE_CBP1_:35Smin_pro_*). The numbers in (a-d) indicating length of promoter were based on *CBP1* promoter taking its transcriptional start site as +1. Left of the boxplots in (a-d) are graphic illustrations of the promoter regions driving the *GUS* reporter gene with the open triangle and grey boxes illustrating the T-DNA insertion site projected onto the *CBP1* promoter and of *CYC-RE_CBP1_*, respectively. Black and blue colour label regions originating from the *L. japonicus* wild-type genome and from the T-DNA in T90, respectively. The larger experimental set-up boxplots including the results of negative controls is presented Fig. S4 with statistical tests. Note the palindromic sequence within the *CYC-RE_CBP1_*highlighted by opposing arrows. Interrupted by only two non-matching basepairs highlighted in grey & bold.

### Quantitative fluorometric GUS assay and analysis

*Nicotiana benthamiana* leaf discs were subjected to a quantitative fluorometric GUS assay (Jefferson 1987) adapted for 96-well plate format. A total number of four to eight leaf discs per indicated combinations was analysed in at least two independent experiments performed in different weeks.

**Genomic DNA extraction and investigation of promoter methylation pattern** is detailed in Method S6.

### Data visualization and statistical analysis

Statistical analyses and data visualization were performed with RStudio 1.1. 383 (RStudio Inc.). Boxplots were used to display data in Fig 2, S3 & S5 (Wickham & Stryjewski, 2011). Individual data points were added to boxplots using R package “beeswarm” (https://github.com/aroneklund/beeswarm). R package “agricolae” (Mendiburu 2017) was used to perform ANOVA statistical analysis with *post hoc* Tukey. Statistical results were presented in small letters where different letters indicate statistical significance, while overlapping letters indicate no significant statistical difference.

## Results

### T90 *white* mutants were identified from an EMS-mutagenised T90 population

To identify the regulators of the T90 *GUS* gene, we performed two independent forward genetic screens. The rationale was as follows: mutants with altered GUS activity (and/or impaired symbiotic behaviour) likely possess defects in the regulatory machinery that directly or indirectly regulate the transcription of the *GUS* gene. Given that transcriptional activation of the *GUS* gene in T90 occurs in response to symbiotic interactions, these impaired machineries potentially regulate gene expression in AM and/or RNS. An EMS-mutagenised T90 M_2_ population was generated by separately harvesting seeds of 1342 M_1_ plants labelled T0001-T1342. Two independent screens were conducted at the seedling stage utilising individual M_2_ families to identify (A) individual M_2_ plants displaying spontaneous activation of the *GUS* gene in the absence of symbionts or (B) individual M_2_ plants with altered GUS activity in presence of *M. loti* (Fig. **2a**; for further details see Tuck 2006). For this purpose, root pieces were removed and stained with 5-bromo-4-chloro-3-indolyl-β-D-glucuronic acid (X-Gluc) whereas the rest of the seedling was maintained to allow for seed production and analysis of heritability. Screen A of 519 M_2_ and 203 M3 lines resulted in 84 plants from 55 lines which showed spontaneous *GUS* expression in the roots, however no progenies from them inherited this phenotype. Screen B of 709 M_2_ lines for loss of *M. loti*-induced GUS activity, resulted in three lines that exhibited heritable aberrant GUS phenotypes. In detail, three M_2_ plants (L8668 and L8686-8687, progeny from M_1_ plant T614 and T1305, respectively), were identified that did not exhibit blue staining after incubation with *M. loti*. Based on the white colour of their roots after GUS staining, these three plants were renamed T90 *white* mutants (L8668 *white1*, L8686 *white2* and L8687 *white3*) and allowed to self-fertilize. The progeny of all three T90 white plants displayed normal shoot and root morphology and could successfully establish AM and RNS similarly as T90 and *L. japonicus* Gifu (Fig. **S2a-b**), however GUS activity could not be detected in their roots during both symbioses (Fig. **2b**, **S2c-d**). T90 *white2* was less healthy than *white3* and produced limited seeds at the time of this study; therefore, only the progeny of T90 *white3* was included in some subsequent experiments.

### T90 *GUS* expression is genetically located downstream of the common symbiosis genes

All three T90 *white* mutants were able to establish both RNS and AM without apparent defects, indicating that essential genes for establishment of symbioses were intact. We sequenced the endogenous *GUS* gene in T90 *white* mutants and did not detect alterations in its coding sequences, hence ruling out that mutations in the endogenous *GUS* genes caused the T90 *white* phenotype. For the cause of T90 *white* phenotype, we considered two possible scenarios: 1) a pathway independent of essential symbiosis genes is involved in activation of the T90 *GUS* gene, which is defective in T90 *white*; 2) the regulatory region of the *GUS* gene is defective in T90 *white* leading to aberrant *GUS* gene induction. To investigate these possible scenarios, we sought to resolve the connection of the transcriptional activation of the T90 *GUS* gene to that of symbiosis signalling by crossing the T90 *GUS* insertion into homozygous backgrounds for mutant alleles of common symbiosis genes. These included mutant allele *ccamk-2* for *CCaMK* [line cac57.3, Schauser *et al.*, 1998, Perry *et al.*, 2009)], *symrk-10* for *SymRK* (Perry *et al.*, 2003), *nfr1-1* for the nod factor receptor gene *Nod Factor receptor 1* (*NFR1*) (Radutoiu *et al.*, 2003) and *pollux-1* for the cation channel *Pollux* (EMS70; Szczyglowski *et al.*, 1998). Preliminary results showed that the F_2_ plants from the following crosses, *ccamk-2 x* T90, *nfr1-1 x* T90 and T90 x *symrk-10* (Gossmann *et al.*, 2012) did not respond with *GUS* expression after *M. loti* inoculation. Based on these findings, we concluded that the T90 *GUS* gene expression is dependent on the tested genes, and positioned the transcriptional activation of the T90 *GUS* gene downstream of *NFR1, SymRK, Pollux* and *CCaMK*. Consistent with this model, ectopically expressed *SymRK* in T90 hairy roots was able to induce the T90 *GUS* expression (Ried *et al.*, 2014).

We tested whether the T90 *GUS* gene expression can be activated by transgenically expressed autoactive CCaMK^T265D^ in T90 and T90 *white* mutants’ hairy roots. These T90 hairy roots spontaneously induced *GUS* expression in the absence of a microsymbiont (Gossmann, 2011).

Importantly, *Agrobacterium rhizogenes*, a member of the Rhizobiaceae and frequently used for hairy root transformation, did not induce the *GUS* expression in T90 (Gossmann, 2011), supporting that the observed spontaneous *GUS* induction resulted from transgenically expressed CCaMK^T265D^. By contrast, we did not observe spontaneous induction of *GUS* expression by CCaMK^T265D^ in T90 *white 1* and T90 *white 3* hairy roots (data not shown). These data provide independent evidence for the position of the T90 *GUS* expression downstream of *CCaMK*. The observation that T90 but not T90 *white* mutants responded with *GUS* expression to autoactive CCaMK^T265D^ combined with the fact that CCaMK phosphorylates and interacts with Cyclops for activation of downstream genes during symbiosis (Singh *et al.*, 2014; Pimprikar *et al.*, 2016; Cerri *et al.*, 2017), motivated us to investigate a possible involvement of Cyclops in the T90 *white* phenotype. We therefore sequenced *Cyclops* in T90 *white* mutants to test for the unlikely case that mutations in the gene prevented *GUS* genes induction, without influencing symbiosis *per se*. We did not detect any sequence alterations of the *Cyclops* gene in all three mutant lines.

### Transgenic insertion of a T90 promoter:GUS fusion in the T90 *white* mutant background restored symbiosis-inducible *GUS* expression

Based on the observed dependency of T90 *GUS* expression on genes involved in early symbiotic signalling, together with the success of T90 *white* to form both RNS and AM, we hypothesized that these symbiosis genes are likely functional in T90 *white*. We consequently redirected our focus onto the regulatory region of the T90 *GUS* gene. To this end we cloned a chimeric region of 2530 bp directly 5’ of the *GUS* gene in T90, hereafter called “T90 promoter”. This region comprised a 1942 bp fragment positioned between −2870 bp to −929 bp relative to the transcriptional start site of *CBP1*, followed at the 3’ end by 588 bp of the T-DNA sequence 5’ of the ATG of the *GUS* gene (Fig. **3a**). This fusion is identical to the original T90 fusion and contains all elements necessary for the transcription of the *GUS* gene, such as a minimal promoter and a transcriptional start site (Jefferson *et al.*, 1987; Topping *et al.*, 1991). We transformed *L. japonicus* Gifu hairy roots with T-DNAs containing a *GUS* reporter gene driven by the T90 promoter (*T90_pro_:GUS*) and analysed the *GUS* expression in the transgenic roots followed by inoculation with *M. loti* MAFF 303099 *Ds*Red (*M. loti Ds*Red). Upon exposure of roots to X-Gluc, blue staining indicative of *GUS* expression was detected exclusively in root hairs (Fig. **S3c**) and nodules (Fig. **3a**,**S3a**). By contrast, *T90_pro_:GUS*-transformed roots grown in the absence of microsymbionts or roots transformed with an identical construct in which the T90 promoter was replaced by a 4 bp spacer sequence did not exhibit any blue staining (Fig. **3a**,**S3a**). The staining pattern achieved by the *T90_pro_:GUS* fusion in transgenic roots resembled strongly the pattern observed in T90 completely matching each other in all key aspects investigated: expression in response to *M. loti* inoculation exclusively in root hairs in early infection stage and later in nodules, which eventually disappeared in mature nodules.

We sequenced the corresponding T90 promoter region of the three T90 *white* mutants and could not detect any sequence alteration (data not shown). We consequently hypothesised that these mutants may suffer from an epigenetic change that renders its corresponding T90 promoter region non-functional. To test this hypothesis, T90 *white1* and *white3* hairy roots were transformed with the *T90_pro_:GUS* reporter fusion construct which gave nodulation-specific *GUS* expression in *L. japonicus* Gifu hairy roots and analysed the *GUS* expression after inoculation with *M. loti Ds* Red. Because the endogenous *GUS* genes in the T90 *white* mutants were not induced during nodulation, any detected blue staining could only result from expression of the introduced reporter gene. Blue staining could be observed in root hairs (Fig. **S3c**) and nodules (Fig. **3b**,**S3b**) in transformed roots of both T90 *white1* and *white3*. By contrast, T90 *white1* hairy roots transformed with a *GUS* reporter gene driven a 4 bp spacer sequence did not exhibit any blue staining (Fig. **3b**,**S3b**). When these transgenic roots were grown in the absence of microsymbionts, no blue staining was observed. These observations suggested that the machinery targeting the T90 promoter to induce gene expression during nodulation is intact in T90 *white* mutants and supported the hypothesis that epigenetic changes block the expression of the endogenous *GUS* genes in T90 *white* mutants.

### A 54 bp and a 113 bp region in the T90 promoter are required for tissue specific expression

To further dissect the promoter and identify relevant regions and *cis*-elements, we generated 5’ deletion series of the T90 promoter in the context of the *T90_pro_:GUS* reporter fusion (starting at −2870 bp relative to the *CBP1* transcriptional start site), without modifications of the rest of the promoter or reporter. The resulting individual constructs started at −1327, −1146, −1092 and −979 bp (Fig. **2c**) and were introduced individually into *L. japonicus* Gifu or T90 *white1* hairy roots. Blue staining was observed in nodules transformed with either of the T90 promoter:GUS fusions starting at −1327, −1146 or −1092 bp (Fig. **3b**). Further deletion to −979 bp, as well as promoter replacement with a 4 bp spacer sequence eliminated blue staining in nodules (Fig. **3b**). An identical pattern was observed in hairy roots of T90 *white1* where a −1092 bp region could achieve GUS activity in transgenic nodules, but not a −979 bp region (Fig. **3c).** We also observed blue staining in patches of root hairs of Gifu and T90 *white* roots transformed with the *GUS* reporter gene driven by *T90_pro_* or a shorter promoter −1146 bp (Fig. **S4c**), fully resembling the T90 *GUS* pattern in the root epidermis (Fig. **1b**,**S1**). The epidermal blue staining was no longer detected when a region −1092 bp was tested (Fig. **S4c**). This 54 bp region (−1146 to −1092 bp) was therefore called the “*Root Hair Element in the CBP1 promoter*” (*RHE_CBP1_*). We concluded based on these findings that a region of 113 bp (−1092 to −979 bp) and a stretch of 54 bp located directly 5’ (−1146 to −1092 bp; *RHE_CBP1_*) was necessary for gene expression in nodules and the root epidermis, respectively.

### T90 promoter hypermethylation was detected in three T90 *white* mutants

DNA methylation is an important and frequently occurring driver of epigenetic changes, which can attenuate binding by the transcription regulatory proteins, thereby inhibiting the activation of target genes (Medvedeva *et al.*, 2014; Yang *et al.*, 2020). We investigated whether an epigenetic event interfering with the endogenous *GUS* expression in T90 *white* mutants could be related to DNA methylation. To detect differences in the methylation pattern between T90 and T90 *white* mutants, we took advantage of restriction digestion enzymes, whose endonucleolytic activities are differentially affected by DNA methylation patterns in their recognition sites. We focused on the short 113 bp region because deletion of this region led to a complete absence of *GUS* gene expression during nodulation. We performed restriction digestion of genomic DNA (gDNA) followed by Polymerase Chain Reaction (PCR). Four enzymes - *Hae*III, *Apa*LI and *Ava*II that are sensitive to DNA methylation and *Bpi*I that is insensitive were chosen because their recognition sites were within or directly adjacent to this 113 bp region (Fig. **4a**). Unlike *Bpi*I that cleaves the target DNA regardless of the methylation pattern, endonucleolytic cleavage by *Hae*III, *Apa*LI or *Ava*II is blocked by a specific methylation signature present at their recognition sites (https://rebase.neb.com). Consequently, amplification of the DNA region of interest in a PCR using gDNA digested with *Hae*III, *Apa*LI or *Ava*II as a template is only successful when their recognition sites were methylated. We subjected gDNA extracted from roots of *L. japonicus* Gifu, T90 and T90 *white* mutants grown in the absence of microsymbionts to restriction digestion by either one of these enzymes. A promoter region of the *NIN* gene that did not contain any recognition site could be successfully amplified with digested gDNA from all genotypes, demonstrating that the gDNA quality was suitable for PCR after restriction digestion (Fig. **4b;** bottom panel). When the gDNA was digested with *Hae*III, *Apa*LI or *Ava*II, a PCR product corresponding to *CBP1* promoter was obtained using digested gDNA from T90 *white* mutants as the amplification template, but not with that from Gifu or T90 (Fig. **4b;** top panel). In contrast, no PCR product was obtained with *BpiI-*digested gDNA regardless of the genotype. Altogether, these results indicated that all three T90 *white* mutants were hypermethylated within and around this 113 bp region (Fig. **3c**).

### *CBP1* is regulated by the CCaMK/Cyclops complex

The observations that the T90 *GUS* gene expression can be induced by exposure to rhizobia and an AM fungus as well as spontaneously by autoactive CCaMK suggested that the T90 promoter is likely subject to CCaMK/Cyclops regulation. We used transient expression assays in *Nicotiana benthamiana* leaves to test whether the CCaMK/Cyclops protein complex could transcriptionally induce expression of a *GUS* reporter gene under the control of the *CBP1* promoter or the T90 promoter (Fig. **5a**, **S4a-b**). A 2870 bp region 5’ of the transcriptional start site of *CBP1* was cloned together with the 177 bp 5’ untranslated region (UTR) of *CBP1* from *L. japonicus* Gifu (*CBP1_-2870pro_*). The expression of the reporter gene driven by *T90_pro_* or *CBP1_−2870pro_* was induced in the presence of Cyclops and the autoactive CCaMK^1-314^ (CCaMK^1-314^/Cyclops; Fig. **5a**,**S4b**). In addition, *T90_pro_* achieved transcriptional activation mediated by CCaMK^T265D^/Cyclops (Fig. **S4a**). A 928 bp stretch of *CBP1* promoter corresponding to the region 3’ to the T90 T-DNA insertion site (*CBP1_-928pro_*; fused to the *CBP1* 5’ UTR in the reporter fusion) did not achieve reporter gene induction by CCaMK^1-314^/Cyclops (Fig. **5a**,**S4b**). These observations together indicated the presence of putative *cis-*regulatory elements responsive to CCaMK/Cyclops-mediated transactivation between −2870 and −928 bp of the *CBP1* promoter.

### *CBP1* promoter dissection revealed a novel Cyclops response *cis*-regulatory element

To identify the CCaMK/Cyclops-responsive *cis-*regulatory element, we generated a promoter 5’ deletion series and investigated reporter gene activation by CCaMK^1-314^/Cyclops-mediated transactivation in transient expression assays in *N. benthamiana* leaves (Fig. **5b**-**d**,**S4c-e**). The 5’ deletion series was built on the basis of *T90_pro_:GUS* (constructed in the same way as those tested in Fig. 3). Each construct comprises a *CBP1* promoter stretch of variable length. The nucleotide position at the 5’ end of the deletions is based on the coordinates of the *CBP1* promoter (see Fig. **1a**). An initial comparison of the transactivation strength across the deletion series revealed a reduction to approximately 50% when comparing −2780 and −2365 bp and a complete loss of activity when comparing −1327 to −979 bp (Fig. **5b**), indicating that these two regions might contain the responsible *cis*-elements. Testing further deletions that were ca. 100 bp different in length within these two regions revealed that the series between −2870 and −2365 exhibited large variations in responsiveness between replicates and was therefore not investigated further. In the −1327 to −979 series, fragments equal to or longer as −1092 resulted in similar reporter gene activation mediated by CCaMK^1-314^/Cyclops, while construct −979 was inactive, suggesting the presence of relevant *cis-*element(s) between −1092 and −979 bp (Fig. **5c**,**S4c**). By testing a higher resolution series with 35 bp to 50 bp length difference, we narrowed down the responsible *cis*-element to the 30 nucleotides between −997 and −967 bp that contained an almost perfect (only two non-matching basepairs; Fig. **5e**) palindromic sequence of 16 bp. We called this element “*Cyclops-response element within the CBP1 promoter*” (*CYC-RE_CBP1_*) because a loss of reporter gene induction by CCaMK^314^/Cyclops was observed when this element was deleted (Fig. **5d**,**S4d**). To test the relevance of *CYC-RE_CBP1_* in the context of the T90 promoter, we mutated or deleted *CYC-RE _CBP1_*(*T90_pro_::mCYC-RE_CBP1_* and *T90_pro_::∆CYC-RE_CBP1_*, respectively). Both resulted in an almost complete loss of CCaMK^1-314^/Cyclops-mediated transcriptional activation, indicating that *CYC-RE_CBP1_* was essential for this transcriptional activation (Fig. **5e**,**S4e**). Moreover, *CYC-RE_CBP1_* fused to a 35S minimal promoter (*CYC-RE_CBP1_:35Smin_pro_*) was sufficient for the activation of reporter gene (Fig. 5e,**S4e**). These results together indicated that *CBP1_pro_*(and *T90_pro_* in the context of T90 genome) is regulated by the CCaMK/Cyclops complex through a *cis*-regulatory element *CYC-RE_CBP1_*.

### *CYC-RE_CBP1_* drives gene expression during RNS and AM

*CYC-RE_CBP1_* is located only 39 bp 5’ of the T-DNA insertion site in T90 and we noticed that *CYC-RE_CBP1_* sits within the hypermethylated region in the T90 *white* mutants (Fig. **3**). Given the necessity and sufficiency of *CYC-RE_CBP1_* for CCaMK/Cyclops-mediated transcriptional activation (Fig. **5**) as well as the common requirement of this protein complex in AM and RNS, we hypothesised that this *cis-* element might be responsible for the symbioses-specific *GUS* expression in T90. To test this, a *GUS* or *DoGUS* (a variant of *GUS*) gene driven by *CYC-RE_CBP1_* fused to a 35S minimal promoter (*CYC-RE_CBP1_:35Smin_pro_*) or the T90 promoter (*T90_pro_*) was introduced into *L. japonicus* Gifu hairy roots, followed by inoculation with *M. loti Ds* Red or the AM fungus *Rhizophagus irregularis (*Fig. **6**,**S3e**&**d**). During nodulation, GUS activity in roots transformed with *CYC-RE_CBP1_:35Smin_pro_:GUS* exhibited blue staining specifically in nodule primordia and nodules, but not root hairs (Fig. **6a**). The same promoter:reporter fusions constructed with *DoGUS* instead of *GUS* led to similar results (Fig. **S3d**). During mycorrhization, blue staining were detected in segments in roots transformed with *T90_pro_:DoGUS* and correlated strongly with the presence of *R. irregularis*, at the entry site of fungal hyphae crossing the epidermis (Fig. **S3e**) and in cortical cells containing arbuscules (Fig. 6b). Blue staining in roots transformed with *CYC-RE_CBP1_:35Smin_pro_:DoGUS* could be specifically detected in the cortex in segments of roots, where cells were infected by *R. irregularis* (Fig. **6b**). In both cases, GUS activity was visibly stronger in cells that were just invaded or had developing arbuscles, compared to those that the arbuscles almost occupying the entire cells. By contrast, roots transformed with *GUS* or *DoGUS* driven by the 35S minimal promoter did not display GUS activity during RNS or AM. Roots transformed with either one of the mentioned fusion constructs grown in the absence of microsymbionts exhibited only rarely blue staining, and if so, in vasculature or root tips regardless of the reporter fusion. We concluded that *CYC-RE_CBP1_* confers AM- and RNS-related gene expression specifically in the fungal-colonised root cortical cells and in nodules, respectively.

**Fig. 6.**
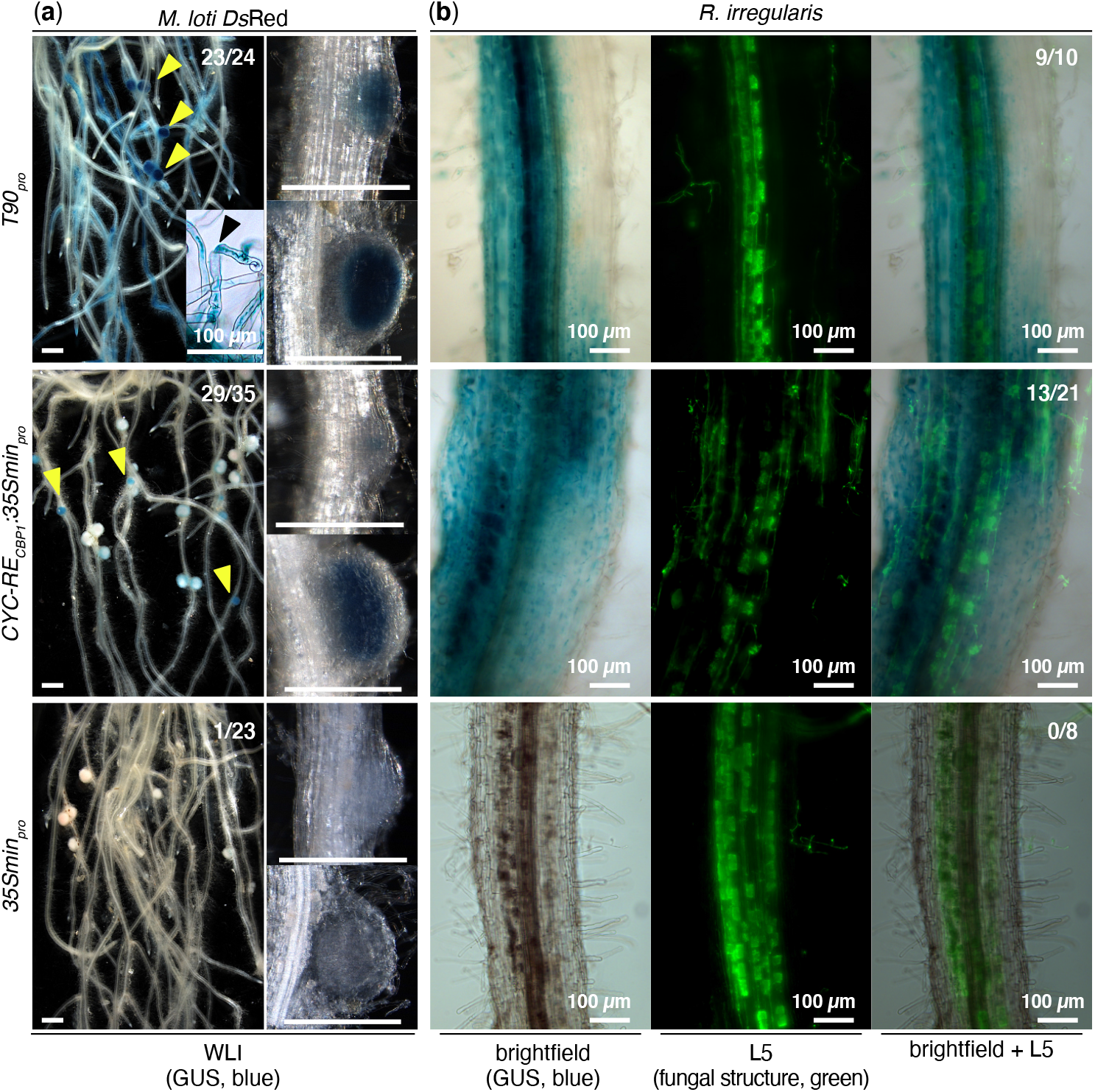
Spatio-temporal *GUS* expression driven by *CYC-RE_CBP1_* in *L. japonicus* hairy roots during nodulation and mycorrhization. *L. japonicus* Gifu hairy roots were transformed with T-DNAs carrying a *Ubq10_pro_:NLS-GFP* transformation marker together with a *GUS* reporter gene driven by the either of the following promoters: the T90 promoter (*T90_pro_*), a 35S minimal promoter (*35Smin_pro_*), or *CYC-RE_CBP1_* fused to *35Smin_pro_*(*CYC-RE_CBP1_:35Smin_pro_*). Transformed roots and nodules were analysed (a) 10 - 14 dpi with *M. loti Ds* Red or (b) 12 dpi with *R. irregularis*. Note that GUS activity was detected in root hairs (black arrowhead in a) in the roots transformed with *T90_pro_:GUS* but not with *CYC-RE_CBP1_:35Smin_pro_:GUS* (for additional support see Fig. S3e). Overall, *T90_pro_:GUS* gave stronger GUS activity (i.e. darker blue colour) in nodules than *CYC-RE_CBP1_:35Smin_pro_:GUS* (yellow arrowheads; compare the overview images of root systems). Note that *35Smin_pro_:GUS* did not show any GUS activity during nodulation or mycorrhization except for GUS activity in vasculature in rare cases. Green: Alexa Fluor-488 WGA-stained *R. irregularis* visualised with a Leica Filter Cube L5. #/#, number of plants showing GUS activity in nodules or root cortex / total number of transgenic root systems analysed. Bars, 1 mm unless stated otherwise.

### The *CBP1* promoter drives gene expression during RNS

We observed that *CYC-RE_CBP1_* in the context of the T90 promoter mediated responsiveness to transactivation by CCaMK/Cyclops and conferred gene expression during symbioses. The T-DNA insertion in T90 physically separated the promoter of *CBP1* into two regions: one containing *CYC-RE_CBP1_* located 5’ of the insertion (5’ region) and the other 3’ of the insertion (3’ region). It has been hypothesized previously that the 5’ region enhances *CBP1* expression during symbiosis while the 3’ region was responsible for its basal expression (Tuck, 2006). To investigate the role of the 3’ region in more detail, we generated *L. japonicus* Gifu hairy roots transformed with a *GUS* reporter gene driven by *CBP1_-2870pro_* that represents the “native” full length promoter comprising both the 5’ and the 3’ region or *CBP1_-928pro_* consists of only the 3’ region (Fig. **1b**,**4a**). Transgenic roots were analysed 14 or 21 dpi with *M. loti Ds* Red for *GUS* expression (Fig. **S5**). *CBP1_-2870pro_:GUS*-transformed roots exhibited strong blue staining in nodules, vasculature tissue, lateral root primordia and root tips (93% of transgenic root systems displaying blue staining in nodules). In comparison, blue staining in *CBP1_-928pro_:GUS*-transformed roots was observed in the same tissue and organ types, however at a lower efficiency (ca. 50% of transgenic root systems displaying blue staining in nodules), and the blue staining was overall visibly weaker in nodules. These observations were consistent with the hypothesis that the 5’ region enhances *CBP1* expression during nodulation.

## Discussion

### The CBP1 promoter comprises at least 4 expression-modulating regions

With the goal to uncover regulatory elements involved in symbiosis, we investigated regulatory circuits underlying the symbiosis-specific *GUS* expression of the *L. japonicus* promoter tagging line T90. We observed that the T90 *GUS* expression can be largely recapitulated in hairy roots transformed with a *GUS* reporter gene driven by a region between −2870 and −967 bp of the *CBP1* promoter fused to the same T-DNA border found in the genomic arrangement of the T90 line. The expression pattern achieved by this region matches all key aspects of that of the T90 line: in root hairs and nodules in presence of *M. loti*; as well as root epidermis and cortical cells when roots were colonised by an AM fungus; and in both symbioses, absent from other tissues such as root vasculature and root tips. We therefore used this transgenic setting as the starting point to dissect the promoter function using a classical promoter deletion series. Our analysis revealed at least four regions/elements with significant impact on *CBP1* expression (Fig. **7**).

**Fig. 7.**
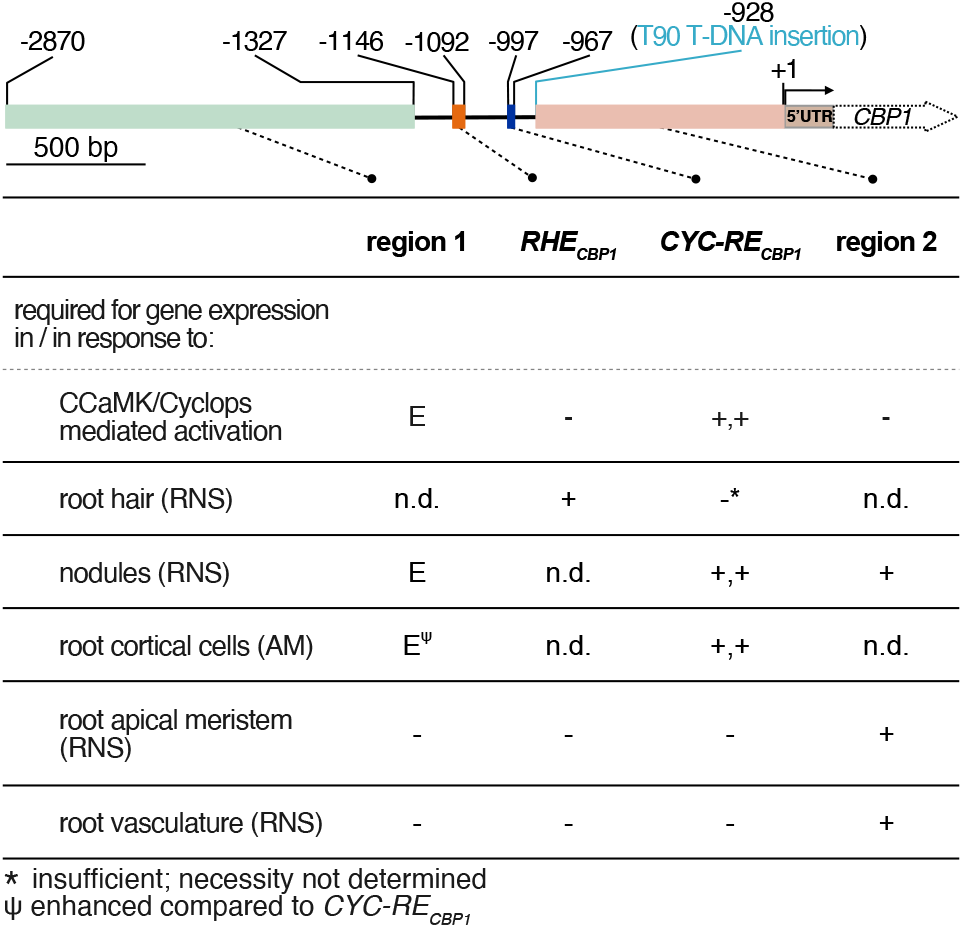
Four regulatory regions of the *CBP1* promoter and their impact on gene expression. *RHE_CBP1_* and *CYC-RE_CBP1_* confer tissue specificity while region 1 and region 2 contribute to the expression strength of the *CBP1* gene. +: necessary; - : not necessary; E: enhancing expression strength; +,+: necessary and sufficient; n.d. : not determined. Note that the quality attribute “necessary” is based on results obtained in different promoter contexts. For *RHE_CBP1_* and *CYC-RE_CBP1_*, this annotation is based on the T90 promoter context and region 2 may contain at least partially redundant functions. The dotted arrow indicating it is not drawn to scale.

#### 1. *A 30 bp CYC-RE*_*CBP1*_ is essential for gene expression in nodules and root cortex

We identified a 30-nucleotide long element named *CYC-RE_CPB1_* within the region −997 and −967 bp which is only 39 bp 5’ of the T-DNA insertion in T90 (Fig. 7). This element, when equipped with a minimal promoter, was able to confer gene expression during both RNS and AM (Fig. **6**), specifically in nodules and infected cortical cells, respectively. The features of *CYC-RE_CBP1_* provide a plausible explanation for the common and symbioses-specific GUS activity in T90: as a result of the T-DNA insertion in T90, the promoterless *GUS* gene was coincidently brought in proximity at the 3’ of *CYC-RE_CBP1_*, a *cis*-element that drives gene expression during colonisation by rhizobia and AM fungi. In the presence of microsymbionts, the *GUS* gene was consequently activated generating a symbioses-specific expression pattern. Our results provide evidence of the involvement of CCaMK/Cyclops in mediating activation of the *CBP1* gene encoding a putative calcium-binding protein for both RNS and AM, through a common *cis-* element *CYC-RE*_CBP1_.

#### 2. A 54 bp region 5’ of CYC-RE_*CBP1*_*(Root Hair Element (RHE*_*CBP1*_)) is essential for gene expression in root hairs

A region between −1146 and −1092 bp was necessary for *GUS* expression in patches of root hairs in proximity to or undergoing IT formation (Fig. **S3c**). Expression in root hairs could not be achieved when the region was deleted from the T90 promoter or when *CYC-RE_CBP1_* was tested on its own.

#### 3. *The region 3’ of the CYC-RE*_*CBP1*_ is boosting CCaMK/Cyclops-mediated expression

The region 3’ of *CYC-RE_CPB1_*, between −928 and −1 (region 2 in Fig. **7**), had on its own very little or no responsiveness to CCaMK/Cyclops mediated gene activation in *N. benthamiana* leaf cells, but significantly boosted such responsiveness in the presence of its 5’ region containing *CYC-RE_CBP1_*(Fig. **5a**). Interestingly, region 2, despite being devoid of a CCaMK/Cyclops response in in *N.* benthamiana leaf cells, conferred gene expression in early nodule development, root vasculature and root tips in *L. japonicus* hairy roots infected by *M. loti*.

#### 4. *The region 5’ of RHE*_*CBP1*_ contains probably multiple regulatory elements

A region between −2870 and −1327 bp (region 1 in Fig. 7) significantly enhances gene activation mediated by CCaMK/Cyclops in *N. benthamiana* leaf cells. Interestingly, the inclusion of this region in *N. benthamiana* transient assays resulted in much larger variation between different leaf discs and unusually strong inter-experimental variation. We interpret this variation as a sensitivity of the underlying regulatory machinery to subtle diurnal, developmental or environmental differences of the leaf tissue that are not observed in other promoter fusions. Deletion of this region did not result in loss of reporter expression in nodules, rather seemed to affect the expression strength of the reporter gene (Fig. **3b**). Hence we focussed only on the reporter gene expression pattern in the roots and did not further investigate a possible quantitative contribution of the −2780 and −1327 bp region.

In summary, we propose a model in which the *CBP1* promoter contains at least 4 distinct regulatory regions that contribute to the expression strength or the tissue specificity or the stimulus specificity of the *CBP1* expression (Fig. **7**).

### The T90 *GUS* expression pattern is governed by the CCaMK/Cyclops complex

By performing transactivation assays in *N. benthamiana* leaves, we observed that *CYC-RE_CBP1_* equipped with a minimal promoter was sufficient for CCaMK/Cyclops-mediated transcriptional activation (Fig. **5e**). Three *CYC-REs* identified earlier come from the promoters of two RNS-induced genes, *NIN* and *ERN1* (*CRE* and *CYC-RE_ERN1_*, respectively; Singh *et al.*, 2014; Cerri *et al.*, 2017) and an AM-induced gene, *RAM1* (*AMCYC-RE;* Pimprikar *et al.*, 2016). The core of *CYC-RE_CBP1_* is semi-palindromic and shares high sequence similarity with *CYC-RE_RAM_1__*and to a lesser extent with *CRE* and *CYC-RE_ERN1_*. Similar to the situation of the three previously identified *CYC-REs*, deletion or mutation of *CYC-RE_CBP1_* drastically impaired Cyclops-mediated transcriptional activation (Fig. **5e**). These observations together suggest that the induction of the *GUS* gene in T90 is at least in the nodules, achieved by *CYC-RE_CBP1_ via* CCaMK/Cyclops-mediated activation and that it contributes essentially, but can not on its own, mediate expression in root hairs (Fig. **6**).

In *L. japonicus*, the expression of *CBP1* during nodulation in roots is likely enhanced by the presence of *CYC-RE_CBP1_* in its promoter (Fig. S5). This conclusion is consistent with the fact that *CBP1* is expressed at a reduced level in T90 roots in which the T-DNA insertion presumably reduces or entirely blocks the activity of this *cis-*element (Webb *et al.*, 2000). Nevertheless, current evidence seems to suggest that the Cyclops-regulated enhancement of *CBP1* expression is dispensable for symbiosis because neither T90 nor T90 *white* mutants are impaired in RNS or AM (Fig. **1**,**2**,**S4**). Given the conserved function of the CaM domain to interact with Ca^2+^ ions, other calcium-binding proteins (e.g., those that identified in Liao *et al.*, 2017) could function redundantly with *CBP1* during symbiosis. Alternatively, it is possible that the residual expression of *CBP1* conferred by the 928 bp regulatory region 3’ of the T-DNA insertion site in T90 (region 2 in Fig. **6**) is sufficient for endosymbiosis or that the impact of the reduced expression levels could not be detected by phenotyping the number of nodules and shoot dry weight of plants (Fig. **S4**).

### The T90 *white* mutant phenotype is caused by cytosine methylation of the T90 promoter

The initial forward genetic approach to screen an EMS-mutagenised T90 population led to the identification of the T90 *white* mutants (Fig. **2**). Based on an analysis using cytosine methylation-sensitive restriction enzymes, we detected hypermethylation within and around in a 113 bp regulatory region of its *GUS* gene (Fig. **4**). This region contains the Cyclops target *cis*-element, *CYC-RE_CBP1_*, which is capable on its own, to drive gene expression during AM and RNS (Fig. **6b**). Cytosine methylation is a well-studied phenomenon and frequently associated with heterochromatin-based gene inactivation (Iwasaki & Paszkowski, 2014). The element’s ability to achieve gene expression during both symbioses and its hypermethylation in the T90 white mutants are in line with the T90 and T90 *white* phenotype. We conclude from all the above that the difference in cytosine methylation in and around *CYC-RE_CBP1_* is the likely cause for the inability the T90 *white* mutants to express their *GUS* genes. In addition, the observation made with T90 *white* mutants indicate that Cyclops activity can be severely impeded by DNA methylation of its target sites. As DNA methylation is overall dynamically altered during nodulation (Satgé, *et al.*, 2016), studying the methylation of *cis-*regulatory elements may reveal another layer of transcriptional regulation during symbiotic development.

### Regulatory elements governing gene activation in both AM and RNS

For most genes that are activated in both RNS and AM, the *cis*-elements responsible for this common induction are not yet known. Only one other element, an AT-rich motif identified in the promoter of *Medicago truncatula ENOD11* gene, was reported important for high-level gene expression during both RNS and AM (Boisson-Dernier *et al.*, 2005). *ENOD11* is one of the earliest marker genes induced by rhizobia as well as an AM fungus (Chabaud *et al.*, 2002; Journet *et al.*, 2001).

The discovery of *CYC-RE_CBP1_* supported a possible scenario that at least a subset of the genes induced during both RNS and AM development could be regulated by the CCaMK/Cyclops complex. This hypothesis is in line with the observations that *CBP1* expression is specifically induced in roots upon Nod factor treatment or inoculation with *M. loti* (data not shown) as well as with the AM fungus *R. irregularis*; and the upregulation of *CBP1* during nodulation is dependent on functional *Cyclops, NFR1* and *NFR5* genes (data retrieved from LotusBase, Webb *et al.*, 2000; Mun *et al.*, 2016). Our observation that four regions of the *CBP1* promoter impact its gene expression suggest that additional pathways are in play to achieve tissue-specific expression pattern and enhance gene expression.

## Acknowledgements

The authors thank Judith K. Webb for the grounding work on the transgenic line T90 and David Chiasson for providing a plasmid containing the *DoGUS* gene that is adapted for the Golden Gate cloning system. E Jensen (née Tuck) thanks Aberystwyth University for a PhD scholarship, which funded the EMS screen. This work was supported by SFB924 “Molecular mechanisms regulating yield and yield stability in plants”. This project has received funding from the European Research Council (ERC) under the European Union’s Seventh Framework Programme (FP7/2007-2013) under grant agreement n° 340904.

## Author Contribution

MP performed mutagenesis of T90 seed to generate the T90 EMS mutant population (Fig. 2a) and EJ conducted screen B. SB performed screen A and investigated the T90 *white* phenotype. XG performed experiments and collected and analysed data (Fig.1, 2b-d, 3 - 5 and all supplementary figures). XG and MP drafted and finalised the manuscript. EJ contributed to manuscript editing.

## Data Availability

All raw data including raw images are either available in the supplement or in repositories. Sequences of constructs and key plasmids necessary for reproducing the results will be made available at Addgene.

