## Supplemental figures and information for "A CCaMK/Cyclops response element in the promoter of *L. japonicus Calcium-Binding Protein 1* (*CBP1*) mediates transcriptional activation in root symbioses"

The following Supporting Information is available for this article:

**Figure S1** T90 roots or nodules displayed GUS activity at 3, 7 or 21 dpi *M. loti Ds*Red.

**Figure S2** T90 *white* mutants lost symbiosis-induced *GUS* expression in the roots but retained symbiosis competence.

**Figure S3** Spatio-temporal *GUS* expression in *L. japonicus* hairy roots transformed with promoter:GUS fusions during nodulation and mycorrhization.

**Figure S4** A *cis*-element in the *CBP1* promoter is necessary and sufficient for CCaMK$^{1-314}$/Cyclops-mediated transactivation.

**Figure S5** *CBP1* promoter-driven reporter gene expression during nodulation in *L. japonicus* roots.

**Table S1** Seedbags used in this study

**Table S2** Constructs and primers used in this study

**Method S1** Plant, bacterial and fungal material

**Method S2** Plant growth condition and phenotypic analysis

**Method S3** Staining method for arbuscular mycorrhizal fungi visualisation

**Method S4** Microscopy

**Method S5** EMS mutant screening

**Method S6** Genomic DNA extraction and investigation of promoter methylation pattern

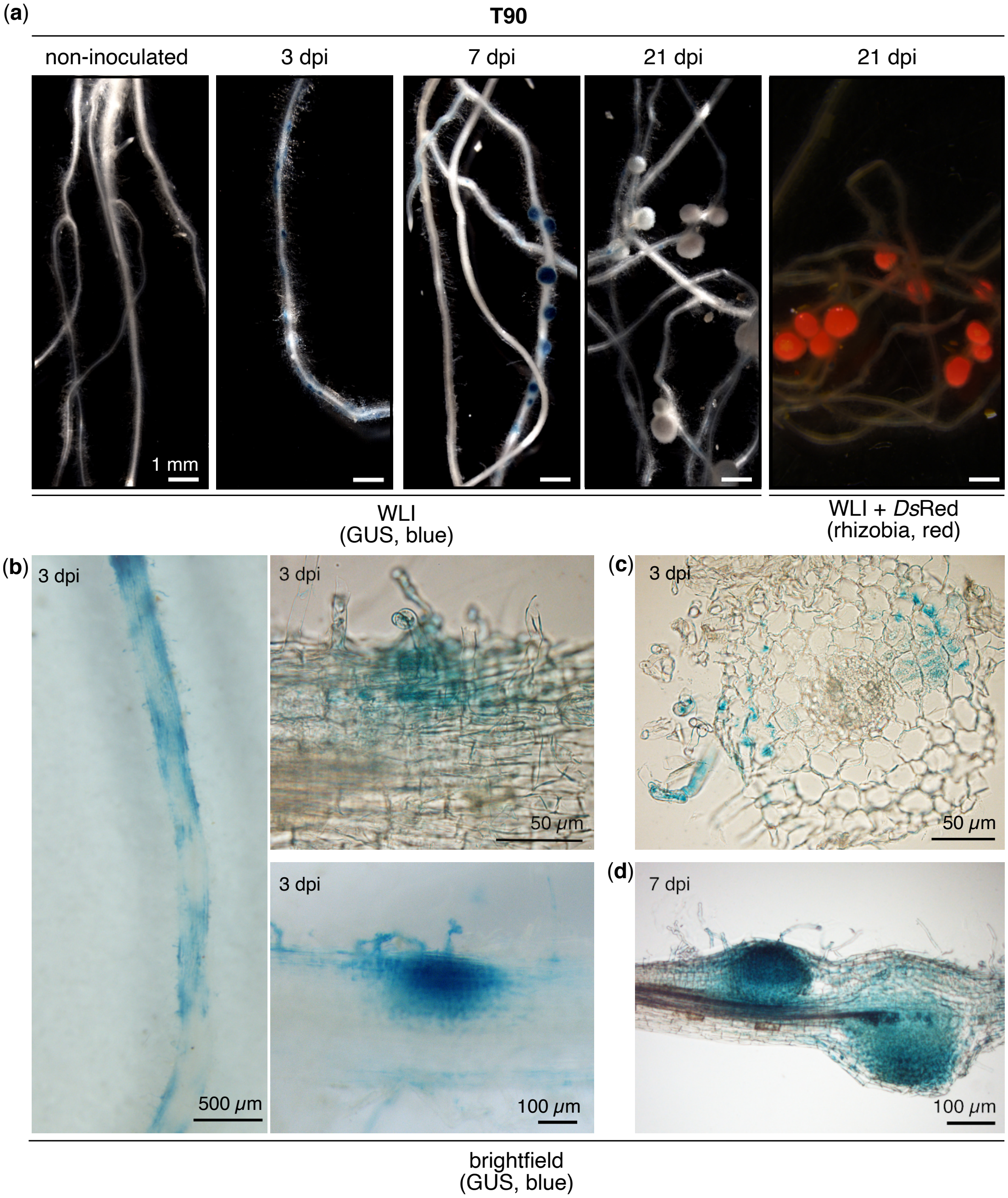


**Fig. S1** T90 roots or nodules displayed GUS activity at 3, 7 or 21 dpi *M. loti Ds*Red. Microscopy images of (a) root systems; (b) magnified areas of roots; and sections of (c) root or (d) nodules. Roots were harvested and stained with X-Gluc at indicated days post inoculation (dpi) with *M. loti Ds*Red. Note that *GUS* expression was induced as early as 3 dpi in root hairs, root epidermis and cortex in (b-c) and later confined into central tissue of nodules at 7 dpi in (d). GUS activity eventually disappeared 21 dpi in mature nodules, which displayed a pink colour characteristic for the leghemolglobin accumulating in legume root nodules in (a). #/# in (a): number of plants displaying GUS activity / total number of plants analysed. Bars, 1 mm unless stated otherwise. WLI: white light illumination.

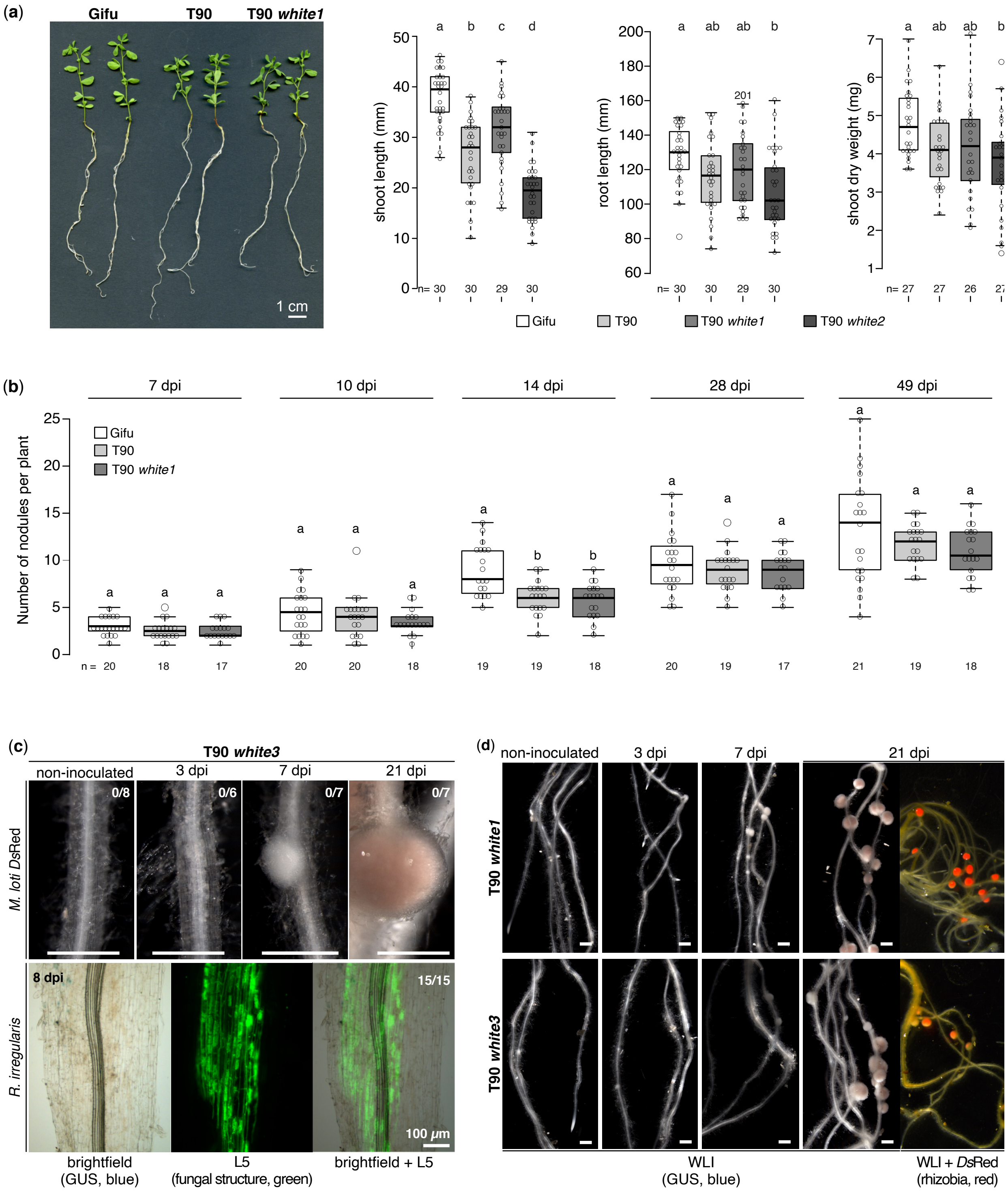
(a)
Gifu
T90
T90 white1
1 cm
shoot length (mm)
root length (mm)
shoot dry weight (mg)
n= 30 30 29 30
n= 30 30 29 30
n= 27 27 26 27
a b c d
a ab ab b
a ab ab b
201
Gifu
T90
T90 white1
T90 white2
(b)
7 dpi
10 dpi
14 dpi
28 dpi
49 dpi
Number of nodules per plant
Gifu
T90
T90 white1
n = 20 18 17 20 20 18 19 19 18 20 19 17 21 19 18
(c)
T90 white3
non-inoculated
3 dpi
7 dpi
21 dpi
0/8
0/6
0/7
0/7
M. loti DsRed
R. irregularis
8 dpi
15/15
100 μm
brightfield
(GUS, blue)
L5
(fungal structure, green)
brightfield + L5
(d)
non-inoculated
3 dpi
7 dpi
21 dpi
T90 white1
T90 white3
WLI
(GUS, blue)
WLI + DsRed
(rhizobia, red)

**Fig S2.** T90 *white* mutants lost symbiosis-induced *GUS* expression in the roots but retained symbiosis competence. (a-b) *L. japonicus* ecotype Gifu, T90 and T90 *white* mutants (T90 *white1* and/or *white2*) were grown in (a) a nitrogen-rich medium (15 mM $KNO_3$) in the absence of symbiont or (b) a nitrogen-poor medium (100 µM $KNO_3$ ) and inoculated with *M. loti DsRed*. Boxplots display (a) the shoot length, root length or shoot dry weight measured 24 days post transfer from the germination medium to the nitrogen-rich medium; or (b) number of nodules quantified at indicated dpi. n: number of plants analysed. (c - d) Roots of T90 *white1* & *white3* were stained with X-Gluc to reveal GUS activity at indicated dpi with *M. loti Ds*Red or AM fungus *Rhizophagus irregularis*. Note the total absence of GUS activity in T90 *white* roots, compared to those of T90 upon inoculation with microsymbionts (tested side-by-side in the same experiment; see Fig. 1a; Fig. S1). Green: Alexa Fluor-488 WGA stained *R. irregularis* visualised with a Leica Filter Cube L5. #/#: number of plants displaying GUS activity / total number of plants analysed. Bars, 1 mm unless stated otherwise. WLI: white light illumination. Statistical method was ANOVA with *post hoc* Tukey: (a) boxplots from left to right, $F_{3,115} = 48.08$, $p < 2\times10^{-16}$; $F_{3,115} = 5.29$, $p = 0.002$; $F_{3,103} = 4.881$, $p = 0.003$; (b) boxplots from left to right $F_{2,52} = 1.418$, $p = 0.251$; $F_{2,55} = 1.514$, $p = 0.229$; $F_{2,53} = 12.49$, $p = 3.6\times10^{-5}$; $F_{2,53} = 0.527$, $p = 0.593$; $F_{2,55} = 1.728$, $p = 0.187$. Different small letters above the boxplots indicate significant difference.

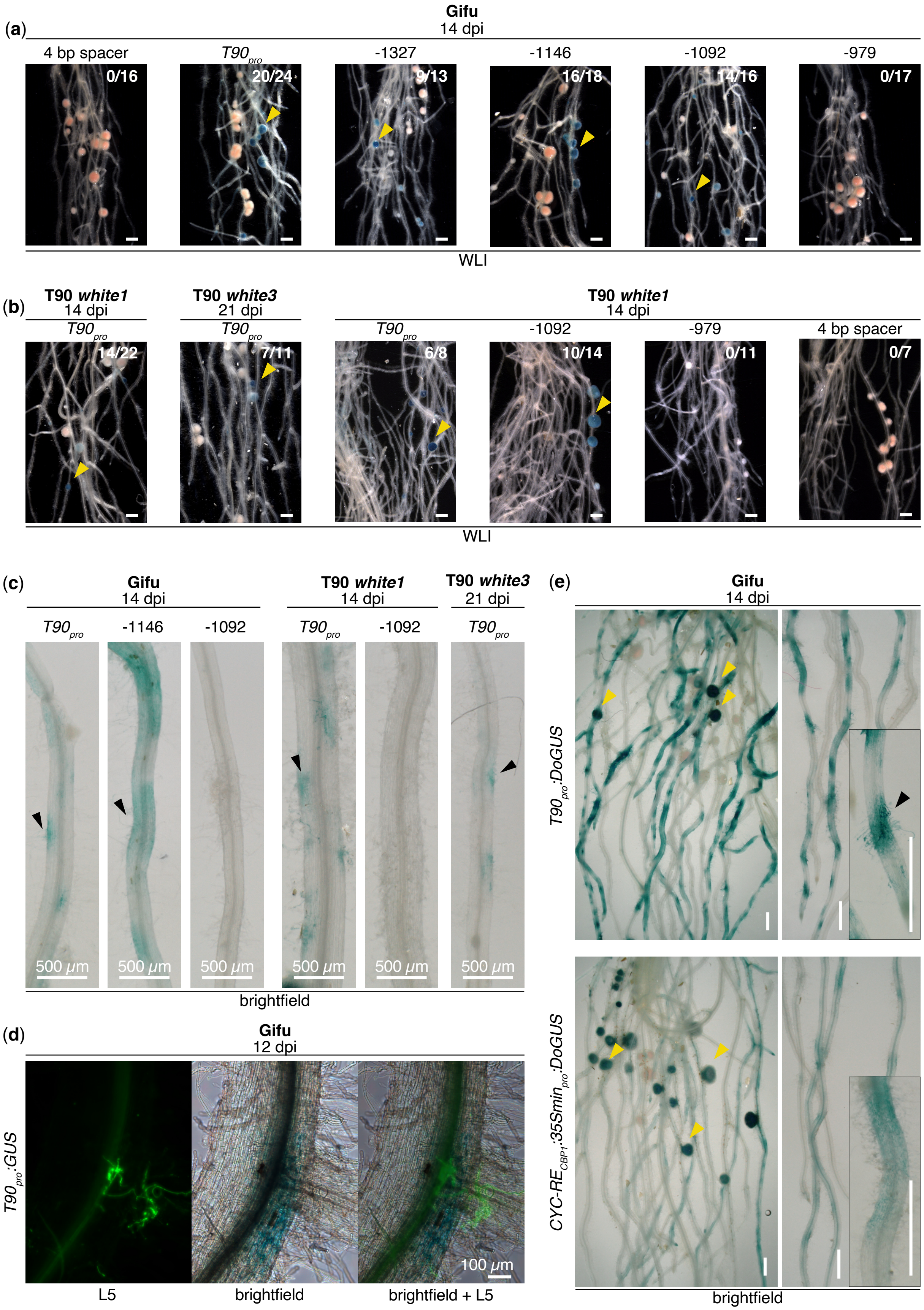

(a)
Gifu
14 dpi
4 bp spacer
T90pro
-1327
-1146
-1092
-979
0/16
20/24
9/13
16/18
14/16
0/17
WLI
(b)
T90 white1
14 dpi
T90 white3
21 dpi
T90 white1
14 dpi
T90pro
T90pro
T90pro
-1092
-979
4 bp spacer
14/22
7/11
6/8
10/14
0/11
0/7
WLI
(c)
Gifu
14 dpi
T90 white1
14 dpi
T90 white3
21 dpi
T90pro
-1146
-1092
T90pro
-1092
T90pro
500 μm
brightfield
(d)
Gifu
12 dpi
T90pro:GUS
100 μm
L5
brightfield
brightfield + L5
(e)
Gifu
14 dpi
T90pro:DoGUS
CYC-RECBP1:35Sminpro:DoGUS
brightfield

**Fig. S3** Spatio-temporal *GUS* expression in *L. japonicus* hairy roots transformed with *promoter:GUS* fusions during nodulation and mycorrhization. (a-b) *L. japonicus* ecotype Gifu or T90 *white* mutant hairy roots transformed with constructs listed in Fig. 3 at indicated dpi with *M. loti Ds*Red; or with *R. irregularis*. The deletion series (starting at -1327, -1146, -1092 and -979 of the *CBP1* promoter) was generated based on the $T90_{pro}$*:GUS* reporter fusion. Note the inability to drive reporter expression in root hairs when the region between -1146 and -1092 bp was deleted (b). GUS activity was lost when more than half of $CYC\text{-}RE_{CBP1}$ was deleted from the promoter region (see -979 bp region in a-b). Note the overall stronger and more widespread GUS activity achieved by the T90 promoter than $CYC\text{-}RE_{CBP1}$*:*$35Smin_{pro}$ (e; see also Fig. 5a). Representative pictures of the nodules from root systems in (a-b) are included in Fig. 3. Black and yellow arrowheads: GUS activity in root hairs and nodules, respectively. #/#, number of plants showing GUS activity in nodules / total number of transgenic root systems analysed. Green in (d-e): Alexa Fluor-488 WGA stained *R. irregularis* visualised with a Leica Filter Cube L5. Bars, 1 mm unless stated otherwise.

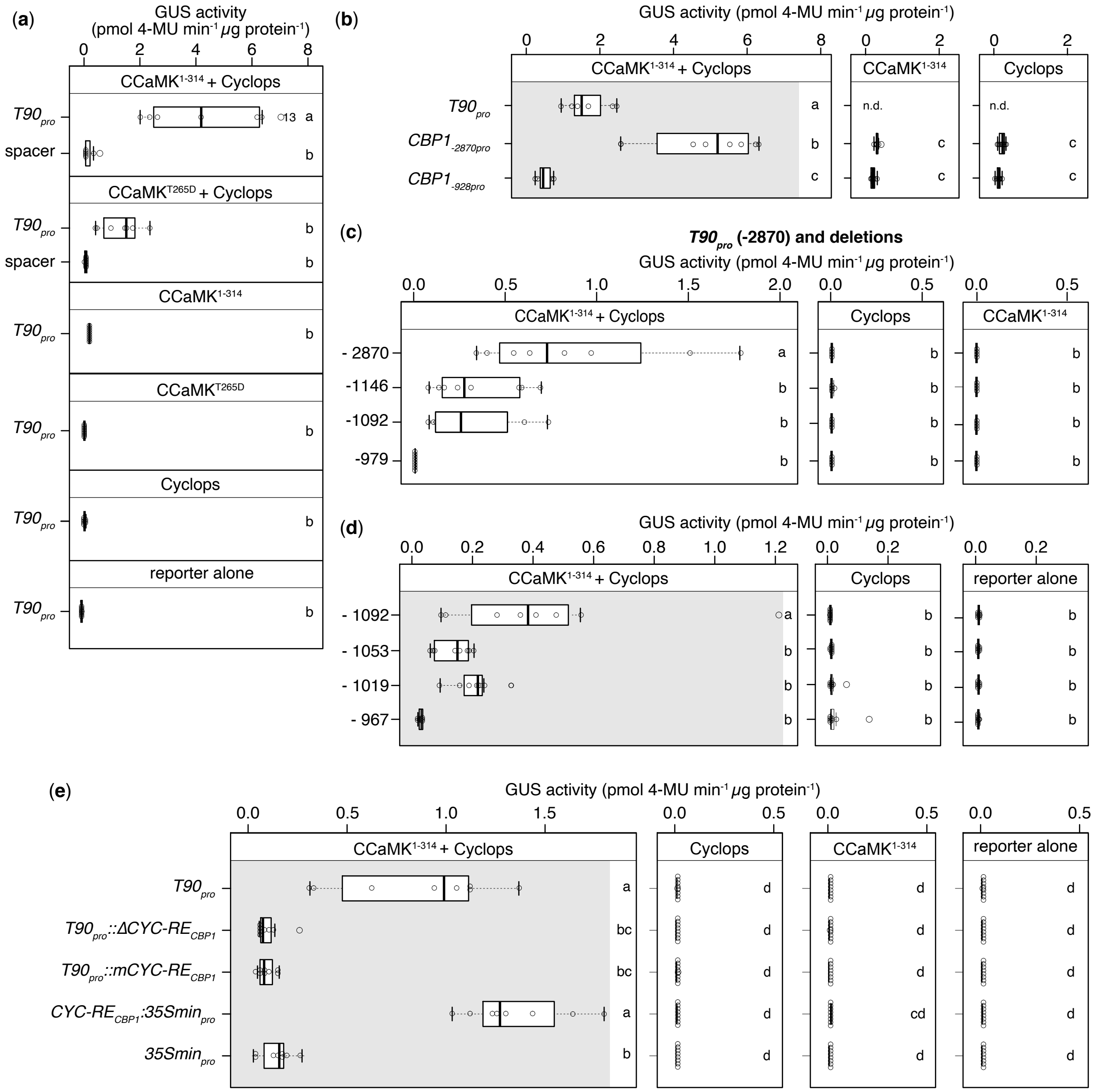


**Fig. S4** A *cis*-element in the *CBP1* promoter is necessary and sufficient for CCaMK$^{1\text{-}314}$/Cyclops -mediated transactivation. *N. benthamiana* leaf cells were transformed with T-DNAs carrying a *GUS* reporter gene driven by either of the indicated promoters: (a) the T90 promoter (labeled as *T90*$_{pro}$ in a-b & e or -2870 in c; see Fig. S3 legend); or a 4 bp spacer sequence; (b) *T90*$_{pro}$; either one of the two *LjCBP1* promoters of varying length (*CBP1*$_{-2870pro}$ or *CBP1*$_{-928pro}$); (c-d) promoter deletion series generated in the context of *T90*$_{pro}$ (see Fig. 5b-d); (e) same constructs as in Fig. 5e. (b-e) include controls for data depicted in Fig. 5a, d & e (grey shaded areas) and the indicated promoter regions in Fig. 5c. The applied statistical method was ANOVA with *post hoc* Tukey: (a), $F_{9,67} = 13.91$, $p = 2.59\times10^{-12}$; (b), $F_{6,47} = 59.55$, $p < 2\times10^{-16}$; (c), $F_{11,52} = 9.558$, $p = 4.05\times10^{-9}$; (d), $F_{11,94} = 40.27$, $p = 7.71\times10^{-9}$; (e), $F_{23,190} = 75.81$, $p = 1.5\times10^{-15}$. Different small letters on the right side of the boxplots indicate significant difference. n.d.: not determined.

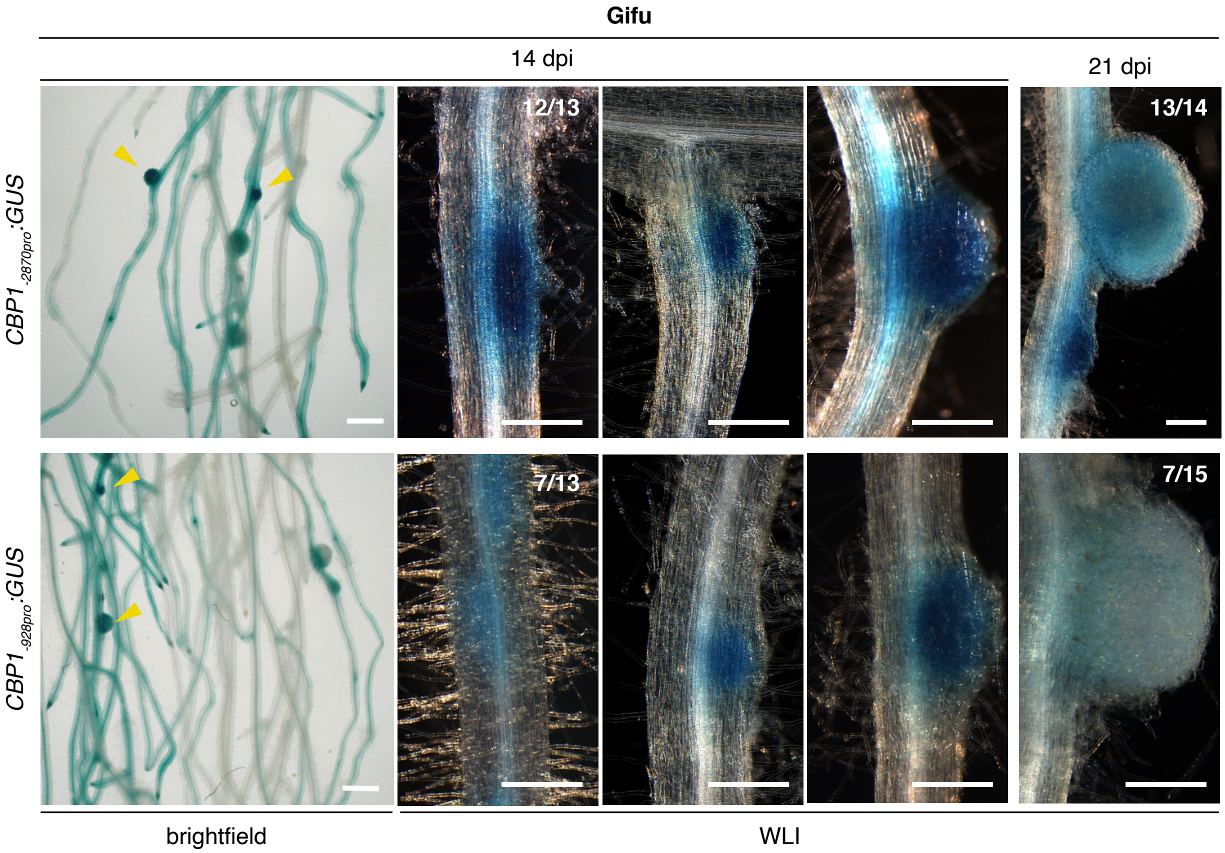


**Fig. S5** *CBP1* promoter-driven reporter gene expression during nodulation in *L. japonicus* roots. *L. japonicus* ecotype Gifu hairy roots transformed with T-DNAs carrying a *Ubq10*$_{pro}$*:NLS-GFP* transformation marker together a *GUS* reporter gene driven by either of the two *CBP1* promoter regions: a 2870 bp region containing *CYC-RE*$_{CBP1}$ (*CBP1*$_{-2870pro}$) or a 928 bp region that did not contain *CYC-RE*$_{CBP1}$ (*CBP1*$_{-928pro}$), were stained with X-Gluc at indicated dpi with *M. loti Ds*Red. Note that only ca. 50% roots transformed with *CBP1*$_{-928pro}$*:GUS* had blue staining in nodules compared to over 93% of those transformed with *CBP1*$_{-2870pro}$*:GUS*. #/#, number of plants showing GUS activity in nodules / total number of transgenic root systems analysed. Bars, 1 mm. WLI: white light illumination.

**Method S1**

Plant, bacterial and fungal material

*Lotus japonicus* genotypes used were Gifu (wild-type, accession B-129, Handberg & Stougaard, 1992); T90 (Webb *et al.* 2000) and the EMS mutant derivatives of T90: T90 *white* 1 (original seeds harvested from plant L8668); T90 *white 2* (original seeds harvested from plant L8686), T90 *white 3* (original seeds harvested from plant L8687). Seed bag numbers are listed in Table S1.

*Mesorhizobium loti* MAFF 303099 constitutively expressing *DsRed* (*M. loti Ds*Red) were used to inoculate *L. japonicus* roots. *M. loti Ds*Red was grown in Tryptone yeast extract (TY) liquid medium (Beringer, 1974) supplied with gentamicin (25 µg/ml) shaken at 180 rpm at 28 °C and harvested by centrifugation at 4000 rpm for 10 min at room temperature (RT). *M. loti Ds*Red was washed twice with FAB medium (500 µM $MgSO_4 \cdot 7H_2O$; 250 µM $KH_2PO_4$; 250 µM KCl; 250 µM $CaCl_2 \cdot 2H_2O$; 100 µM $KNO_3$; 25 µm Fe-EDDHA, catalog no. F0527.0250, Duchefa Biochemie; 50 µM $H_3BO_3$; 25 µM $MnSO_4 \cdot$H2O; 10 µM $ZnSO_4 \cdot 7H_2O$; 0.5 µM $Na_2MoO_4 \cdot 2H_2O$; 0.2 µM $CuSO_4 \cdot 5H_2O$; 0.2 µM $CoCl_2 \cdot 6H_2O$; pH = 5.7), and resuspended in FAB medium to reach a final optical density at 600 nm ($OD_{600}$) of 0.01 for inoculation.

The arbuscular mycorrhizal fungus (AMF) *Rhizophagus irregularis* was used to inoculate *L. japonicus* roots in a chive nurse plant system (based on Wegel *et al.*, 1998). To prepare the nurse plants, *Rhizophagus irregularis* spores (DAOM197198; Connectis, Agronutrition) were used to inoculate chive seedlings (ca. 200 spores for 40 plants). R. irregularis spores were collected by centrifugation at 805 rcf for 10 min at 4 °C and resuspended in 10 ml of 1/4 strength modified Hoagland's solution (based on the nitrogen-free medium described by Hoagland and Arnon in 1938 with the following modifications: 1 mM $KNO_3$ and 100 µM $KH_2PO_4$ added; replacing half of the chelated iron stock solution with 12.5 µm Fe-EDDHA). Chive seeds were briefly sterilised with 1.2 % NaClO and 0.1 % SDS for 1 - 2 min and thoroughly washed with sterile distilled water. Pots used to grow chive plants were washed and sterilised with 70 % ethanol before use. Sterilised chive seeds were placed on the surface of a sterile sand-vermiculite mixture (2:1) in a pot and watered with 35 ml of modified 1/4 Hoagland's solution and spore solution. Chive pots were kept in a growth chamber (24 °C, 16 h light /8 h dark; light intensity of 180 µmol $m^{-2}$ $s^{-1}$) and were covered with a plastic lid for the first 3 days. Chive plants were watered with 1/12 modified Hoagland's solution three times a week (20 ml solution for each pot). Six weeks post inoculation, chive roots were stained to verify AM colonisation. Two chive plants were transferred to a new pot containing a sterile sand-vermiculite mixture and 40 ml of 1/12 modified Hoagland's solution and allowed to grow for another

4 to 6 weeks before being used as nurse plants. The shoot systems of chive nurse plants were cut off for AM inoculation experiments, leaving only colonised roots in the growth substrate.

**Method S2**

Plant growth condition and phenotypic analysis

Seeds were scarified and surface-sterilized as previously described (Groth *et al.* 2010) and plated on ½ Gamborg's B5 with 0.8 % Bacto™ agar plates (Becton, Dickinson and Company). Seeds were kept in the dark in a Panasonic growth chamber 24 °C for three days and then on a 16h light /8 h dark cycle. For phenotypic analysis under symbiotic condition (Fig. **1,2,S1,S2b-d**), 10-day-old seedlings were transferred to Weck jars (SKU745; J. WECK GmbH u. Co. KG) containing 300 ml of sterile sand:vermiculite mixture (2:1) wet with 30 ml of FAB medium containing *M. loti Ds*Red. Roots were harvested indicated days post inoculation stated in the figure legends (dpi) and the number of nodules was quantified. Roots harvested 3, 7 and 21 dpi were subjected to GUS staining as described in Groth *et al.* (2010) with the incubation time of 6 h (Fig. **1,2,S2c-d**) to detect GUS activity. For phenotypic analysis under nitrogen-sufficient conditions (Fig. S2a), 7-day-old seedlings were transferred to Weck jars containing 300 ml Seramis:vermiculite mixture (4:1) wet with 30 ml of the nitrogen-containing version of ¼ Hoagland's solution (15 mM $KNO_3$). Plants were harvested at 24 days post transfer. Shoot height was evaluated as the distance between the youngest leaves to the end of the hypocotyl. Root length was measured as the length of the whole root system. while shoot dry weight was measured as after shoots were dried at 60 °C for 1 h in an incubator. For promoter analysis (Fig. **3,6,S3,S5**), plants with transformed roots were transferred to either Weck jars containing 300 ml of sterile sand:vermiculite mixture (2:1) and 60 ml FAB medium containing *M. loti Ds*Red; or chive nursing pots and watered with modified ¼ Hoagland's solution (see Plant, bacterial and fungal material; with $KNO_3$ increased to 9 mM) three times a week. Plants were grown under the same conditions as described for chive plants.

**Method S3**

Staining method for arbuscular mycorrhizal fungi visualisation

To detect AM colocation in roots of *L. japonicus* wild-type, T90 or T90 *white* mutants, plants cocultivated with mycorrhized chive plants were harvested at indicated days post inoculation (transfer to the nurse pots) (Fig. **1,2**,**S2,S3**). Roots or transgenic root systems were subjected to GUS staining for 14 to 16 h at 37 $^{o}$C, followed by cleaning with 50 % ethanol overnight at RT and then cleared with 10 % KOH. Roots were rinsed with water and incubated in 0.1 M HCl for 2 hours at RT before overnight staining with 1 µg ml$^{-1}$ WGA Alexa Fluor 488 (catalog no. W11261; Thermo Fisher Scientific) dissolved in PBS buffer (140 mM NaCl; 2 mM KCl; 10 mM $Na_2HPO_4$; 2 mM $KH_2PO_4$; pH 7.4). Roots were kept in staining buffer at 4 $^{o}$C in the dark until microscopic analysis.

**Method S4**

Microscopy

Pictures of nodules and whole root systems were taken with a Leica MZ16 FA fluorescent stereomicroscope (Fig. **1,2,3,S1,S2,S3a-b,S3d**) or a Keyence VHX6000 digital microscope (Keyence Deutschland GmbH; Fig. **S3c,3e,S5**). Pictures of mycorrhized roots were taken with a Leica DM6B light microscope equipped with a Leica DMC2900 camera (Leica Microsystems). The presence of WGA Alexa Fluor 488-stained AM fungal structures was detected using the Leica Filter cube L5 (512nm - 542nm; size K, catalogue no. 11513880; Leica Microsystems).

**Method S5**

EMS mutant screening

Screening of the T90 EMS population for mutants showing aberrant GUS activity is summarized in section '*EMS mutagenized T90 population and mutant screening*' in Tuck, 2006. To screen for spontaneous GUS activity, $M_2$ seeds were sterilized and imbibed as previously described, and then plated on ½ strength of B&D medium plates with 0.8% agar and final concentration of 1 µM L-α-(2-aminoethoxyvinyl)-glycine (AVG; ethylene biosynthesis inhibitor). Root pieces of *ca.* 1.5 cm long from 7-day-old seedlings were excised and incubated in 300 µl GUS staining solution in a 96-well plate for 18 h at 37 $^{o}$C. Plants with blue coloration in roots were considered putative spontaneous mutants. M3 seeds of these putative mutants were generated and eight $M_3$ individuals per line were tested in the same way to confirm the spontaneous blue phenotype.

**Method S6**

Genomic DNA extraction and investigation of promoter methylation pattern

Roots (ca. 100 mg) from plants grown in the absence of symbiont of each genotype were harvested, frozen and ground in liquid nitrogen with mortar and pestle. Genomic DNA (gDNA) was extracted as previously described (Lueders *et al.* 2004). Concentration of gDNA was determined with a Nanodrop photometer (DS-11; DeNovix Inc.). In total, 25 ng gDNA was subjected to restriction digestion by *Hae*III, *Apa*LI, *Ava*II or *Bpi*I (New England Biolabs GmbH) in a 10 µl reaction that contained 1 µl appreciate NEB restriction buffer (supplied with the enzyme), 1 µl gDNA, 1 µl (ca. 10 units) of enzyme and 7 µl MilliQ water for 18 h at 37 $^{o}$C. PCR was performed with 1 µl of digestion mix as template and the primer pair 5'-AATAGTGGCATATGAAAATGTTGG-3' (F1) and 5'-AATTATAGGAAGACGTTGGAGAGT-3' (R1; Fig. 5) to amplify a 220 bp region in the T90 promoter containing a single recognition site of each of the enzymes, or the primer pair 5'-TTTCGCCGATATCGTAGAC-3' and 5'-GCAACACCGGCTATATAATAGTG-3' to amplify a 199 bp region of the NIN promoter that does not contain recognition sites for any of the enzymes, as a control for the quality of digested gDNA. PCR products were detected using agarose gel electrophoresis (1.5 % agarose gel).
